# Global niche partitioning of purine and pyrimidine cross-feeding among ocean microbes

**DOI:** 10.1101/2024.02.09.579562

**Authors:** Rogier Braakman, Brandon Satinsky, Tyler J. O’Keefe, Krista Longnecker, Shane L. Hogle, Jamie W. Becker, Robert C. Li, Keven Dooley, Aldo Arellano, Melissa C. Kido Soule, Elizabeth B. Kujawinski, Sallie W. Chisholm

**Author notes:** Current address: Dyno Therapeutics, Watertown, MA, USA. Current address: Institute of Marine Sciences, University of North Carolina at Chapel Hill, Morehead City, NC, USA. Current address: Department of Biology, University of Turku, Turku, Finland (Shane L. Hogle). Current address: Department of Science, Alvernia University, Reading, PA, USA. Current address: Renewable Resources and Enabling Sciences Center, National Renewable Energy Laboratory, Golden, CO, USA. Current address: Department of Bacteriology, University of Wisconsin-Madison, Madison, WI, USA.

## Abstract

Cross-feeding involves microbes consuming the exudates of other surrounding microbes, mediating elemental cycling. Characterizing the diversity of cross-feeding pathways in ocean microbes illuminates evolutionary forces driving self-organization of ocean ecosystems. Here, we uncover a purine and pyrimidine cross-feeding network in globally abundant groups. The cyanobacterium *Prochlorococcus* exudes both compound classes, which metabolic reconstructions suggest follows synchronous daily genome replication. Co-occurring heterotrophs differentiate into purine– and pyrimidine-using generalists, or specialists that use compounds for different purposes. The most abundant heterotroph, SAR11, is a specialist that uses purines as sources of energy, carbon and/or nitrogen, with subgroups differentiating along ocean-scale gradients in the supply of energy and nitrogen, in turn producing putative cryptic nitrogen cycles that link many microbes. Finally, in a SAR11 subgroup that dominates where *Prochlorococcus* is abundant, adenine additions to cultures inhibit DNA synthesis, poising cells for replication. We argue this subgroup uses inferred daily adenine pulses from *Prochlorococcus* to synchronize to the daily photosynthate supply from surrounding phytoplankton.

## Introduction

Microbial processes and interactions lie at the heart of the oceanic carbon cycle (1–3). Carbon enters the marine biosphere through photosynthesis (4,5), is transformed through metabolism (6) and viral infections and lysis (7,8), is transferred to higher trophic levels through grazing (2,9), and is ultimately broken back down and released as CO_2_ through bacterial catabolism and respiration (1–3,6). An increasingly recognized link within this global cycle of organic carbon is cross-feeding, in which cells excrete specific compounds that are consumed by surrounding cells (10–13). Evidence that cross-feeding is widespread in the ocean comes from the highly streamlined genomes and complex nutrient dependencies of many highly abundant oceanic bacteria (14–16). However, partly due to the difficulty of measuring extracellular metabolites in seawater (3), the diversity and distribution of cross-feeding pathways has not been systematically characterized, obscuring both the ecological roles they play and the forces shaping their evolution.

To explore this using a bounded model system, we have begun characterizing the production of organic carbon by the highly abundant oceanic cyanobacterium *Prochlorococcus* (17) and assessing its fate. By performing ∼10-15% of oceanic CO_2_-fixation (18,19), *Prochlorococcus* is a major source of organic carbon to ocean ecosystems, and over geologic timescales is thought to have evolved metabolic interdependencies with co-occurring heterotrophs (20). In recent experiments, we noticed that pyrimidines and purines are both abundant exudates in *Prochlorococcus* (21). Here we further examine the production and consumption of these compounds and uncover a globally distributed cross-feeding network involving many abundant oceanic microbes. We begin to characterize the forces driving large-scale niche partitioning within this network.

## Results and Discussion

### Pyrimidine and purine exudation by Prochlorococcus

To better understand organic carbon production by *Prochlorococcus* we measured in a complementary study (21) the extracellular abundance of a large suite of organic compounds in exponentially growing cultures of several strains under a range of light and nutrient conditions. A signal that stood out was that thymidine is among the most abundant extracellular compounds across strains and culture conditions (Fig. 1, Table S1). Here we set out to further study this signal, as the exudation of high levels of this nitrogen-rich pyrimidine is unexpected, since it is generally thought that the genome and proteome of *Prochlorococcus* are highly streamlined to minimize the nitrogen requirements of growth in oligotrophic waters (22,23). We calculated (Methods) that over the course of the experiment production of extracellular thymidine amounted to almost half of what was incorporated into DNA (Fig. 1, Table S1). We also detected adenine and guanine in the exudates of *Prochlorococcus*, and their levels were higher under phosphorus-limitation than in nutrient replete conditions (Fig. 1, Table S1). While we could not precisely quantify these compounds due to an extraction efficiency <1% (21), their uncorrected levels reached ∼2-3% of that of thymidine, suggesting their exudation was at a similar scale (Fig. 1, Table S1).

**Fig. 1.**
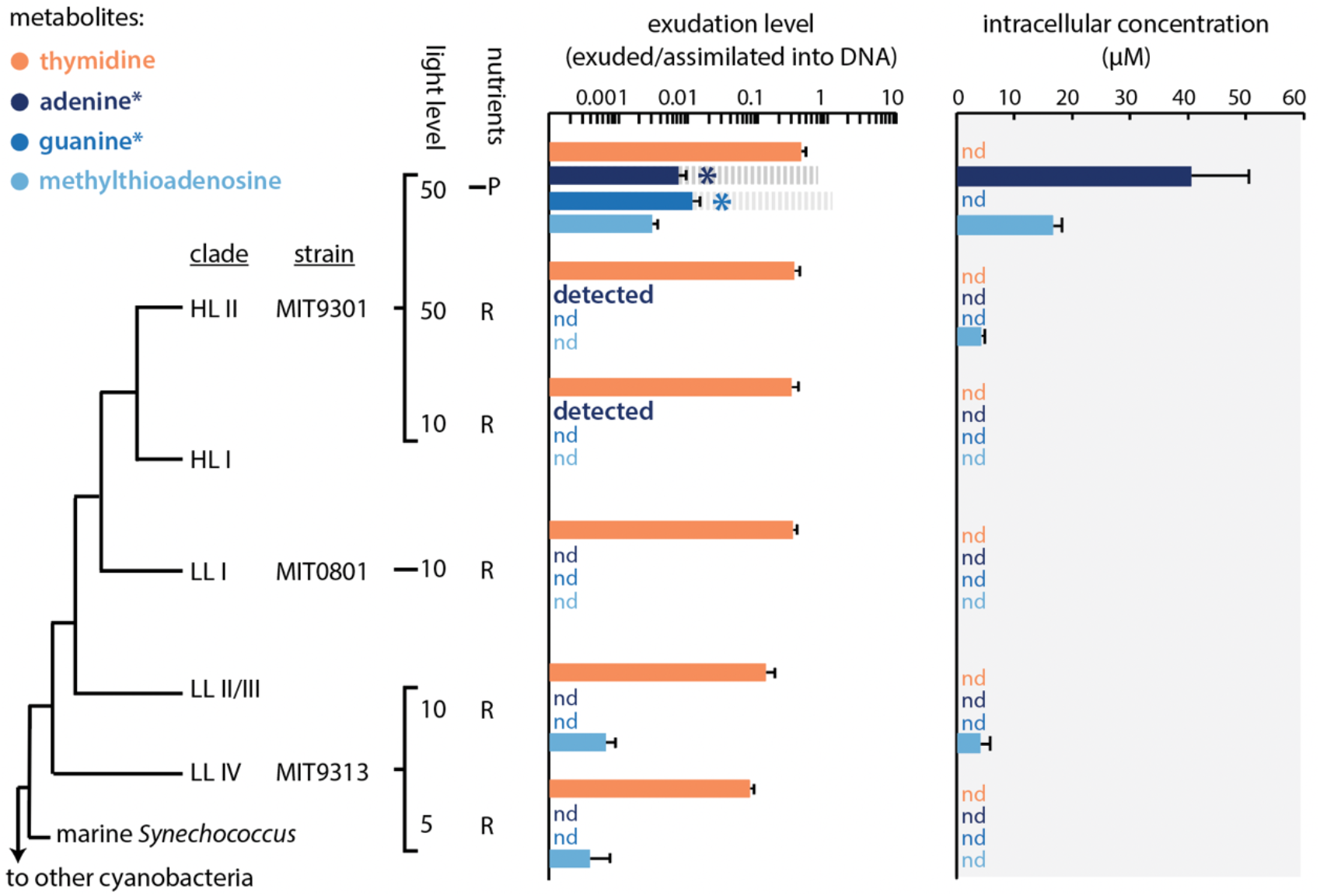
Extra– and intracellular pyrimidines and purines in cultures of *Prochlorococcus*. Extracellular levels of the pyrimidine thymidine (orange), and the purines adenine, guanine and 5-methylthioadenosine (shades of blue) are normalized to the total amounts of purines or pyrimidines incorporated into DNA and shown alongside cytosolic concentrations (gray shading) of these compounds, for three different strains of *Prochlorococcus* grown under different conditions, as measured in a complementary study (21). Error bars show the standard deviation between biological replicates. Light levels are in units of μmol photons m^-2^ s^-1^, ‘R’ stands for batch culture growth in nutrient replete media and ‘–P’ stands for growth under semi-continuous phosphorus limitation. Exudation levels of adenine and guanine (marked with asterisks) are uncorrected from the measured concentrations due to having an extraction efficiency of less than 1%. For a sense of scale relative to thymidine exudation levels, gray dashed bars are included at 100x the measured adenine and guanine levels (i.e. the theoretical correction factor for an extraction efficiency of 1%). Incidences in which metabolites were only detected in single replicates are denoted with ‘detected’, while incidences in which metabolites were not detected in any replicate are denoted with ‘nd’.

To understand the excretion of thymidine, adenine and guanine, we searched for possible pathways producing and/or consuming them in *Prochlorococcus*. The only pathway we identified involving these compounds (Methods, Text S1) is a putative pathway for recycling deoxyribonucleotides, the direct precursors to DNA (Fig. 2A). There thymidine, deoxyadenosine and deoxyguanosine are generated from dTMP, dAMP and dGMP, respectively, through the action of 5’-nucleotidase (SurE) (24). Deoxyadenosine and deoxyguanosine are then further processed to adenine and guanine through the action of methylthioadenosine phosphorylase (MTAP), and then to AMP and GMP through the action of adenine phosphoribosyltransferase (apt), together forming a salvage pathway also observed in other systems (25). We could not identify other genes involved in processing thymidine, suggesting it forms an endpoint and is not further processed after the first (phosphate-salvaging) step of the pathway, perhaps because it is only involved in DNA synthesis, while deoxyadenosine and deoxyguanosine can be repurposed for ATP/GTP and RNA synthesis. This is consistent with the fact that we never measured intracellular thymidine, suggesting it is rapidly excreted after the phosphate is removed from dTMP. Original genome annotations assigned a narrower functionality to MTAP (i.e. acting only on methylthioadenosine) than we hypothesize here, but this enzyme is known to act on multiple substrates in other microbial systems, including as part of the pathway we identify here (25,26). Moreover, enzyme multi-functionality is generally thought to be an important strategy in bacteria with streamlined genomes like *Prochlorococcus* (27,28). We could not identify other pathways involving thymidine, adenine and guanine, nor could we identify other possible roles for SurE, MTAP and apt than in the pathway we reconstructed. Together this leads us to tentatively conclude that thymidine, adenine and guanine are all intermediates or endpoints of a shared, broad specificity deoxyribonucleotide recycling pathway (Fig. 2A).

**Fig. 2.**
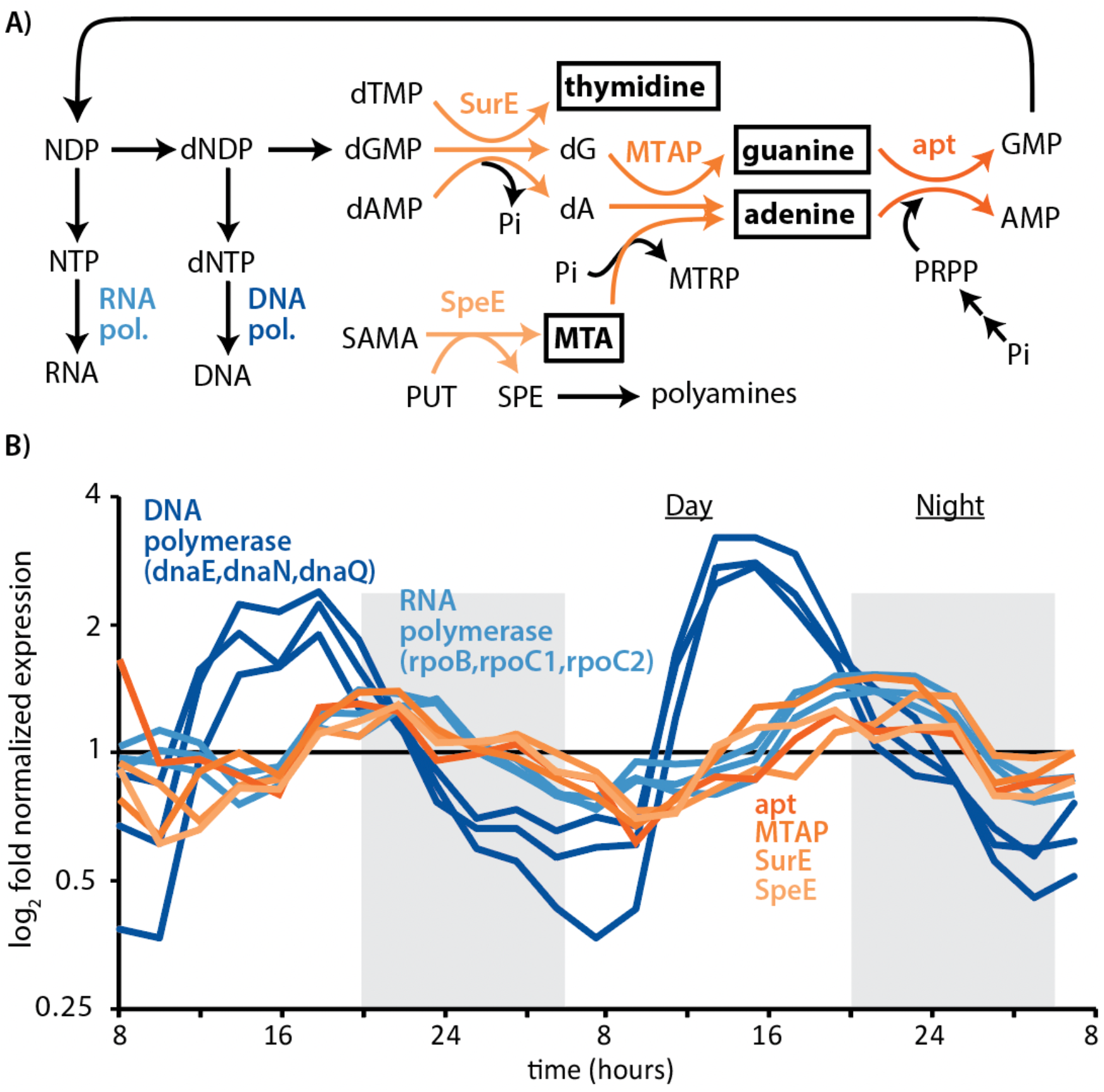
Putative deoxyribonucleotide recycling pathway in *Prochlorococcus*. **A)** Structure of the deoxyribonucleotide recycling pathway in relation to pathways for DNA and RNA synthesis and polyamine metabolism. Metabolites highlighted in boxes are exuded by *Prochlorococcus* (Fig. 1). **B)** transcriptional dynamics of genes involved in the deoxyribonucleotide recycling pathway, as well as subunits of DNA and RNA polymerases, in *Prochlorococcus* MED4 cells synchronized to a diurnal light:dark cycle (29). In both panels deoxyribonucleotide recycling pathway reactions/genes are highlighted using the orange spectrum, and DNA and RNA polymerase are highlighted in shades of blue. Abbreviations: NDP = nucleoside diphosphate, dNDP = deoxynucleoside diphosphate, NTP = nucleoside triphosphate, dNTP = deoxynucleoside triphosphate, dTMP = deoxythymidine monophosphate, dGMP = deoxyguanosine monophosphate, dAMP = deoxyadenosine monophosphate, dA = deoxyadenosine, MTA = 5-methyl-thioadenosine, SAMA = S-adenosylmethioninamine, PUT = putrescine, SPE = spermidine, MTRP = methyl-5-thioribose, PRPP = phosphoribose diphosphate, Pi = orthophosphate, SurE = survival protein E (5’-nucleotidase), MTAP = methylthioadenosine phosphorylase, apt = adenine phosphoribosyltransferase, SpeE = spermidine synthase, dnaE = DNA polymerase III alpha subunit, dnaN = DNA polymerase III subunit beta, dnaQ = polymerase III subunit epsilon, rpoB = RNA polymerase subunit beta, rpoC1 = RNA polymerase subunit betaʹ, rpoC2 = RNA polymerase subunit betaʹʹ.

Since other unknown pathways could still be at play, we examined if genes in the putative deoxyribonucleotide recycling pathway in *Prochlorococcus* exhibit expression, as one might expect if their functions were linked. To this end we analyzed existing transcriptional data obtained from a strain whose growth was synchronized to a diurnal light:dark cycle (29). Under such conditions, metabolic processes are segregated in time, allowing us to examine the potential role of the putative deoxyribonucleotide recycling pathway in the broader processes of growth and cellular replication. The expression of all three genes (SurE, MTAP, apt) in the putative pathway are synchronized and maximally expressed just after dusk, when genome replication is complete and cells transition to RNA synthesis (Fig. 2B), both of which are consistent with their collective functioning in deoxyribonucleotide recycling.

Properties of the deoxyribonucleotide recycling pathway are also consistent with observed metabolite dynamics in *Prochlorococcus* cells under phosphorus-limited growth conditions. That is, regenerating AMP and GMP from adenine and guanine requires phosphate (Fig. 2A), and adenine accumulates intracellularly in phosphate-limited relative to nutrient-replete cells (Fig. 1). This suggests that exudation of adenine and guanine results from their intracellular accumulation and is driven by the emergence of a bottleneck in the deoxyribonucleotide recycling pathway due to a depletion of intracellular phosphate. This is consistent with the observation that methylthioadenosine, a byproduct of polyamine metabolism that is recycled via the same pathway (Fig. 2A), also accumulates and is excreted under phosphate limited relative to replete conditions (Fig. 1). The significantly lower intracellular accumulation of methylthioadenosine compared to adenine (Fig. 1) under similar phosphate-dependent bottlenecks (Fig. 2A), suggests that deoxyribonucleotide recycling, and not polyamine metabolism, is the primary source of adenine in *Prochlorococcus*. A recent study showed that some strains of the purple sulfur bacterium *Rhodopseudomonas palustris* exude adenine when a bottleneck arises in its intracellular salvage due to low levels of the enzyme *apt* (30), highlighting a similar principle. In contrast, the pyrimidine thymidine, which unlike purines is not internally recycled (Fig. 2), never accumulates, and its extracellular levels are not significantly different under phosphorus stress (Fig. 1). Taken together, metabolite, genomic and transcriptional data thus suggest that *Prochlorococcus* cells generate excess dNTP during genome replication and excrete whatever they do not reuse or recycle when transitioning to ribosome synthesis. This in turn implies that *Prochlorococcus* provides daily pulses of pyrimidines and purines to the surrounding ecosystem. Time-resolved quantification of metabolites in the field and in diel L:D synchronized cultures of *Prochlorococcus* are a key area of future study that will help constrain both the timing of release and the relative contributions of *Prochlorococcus* to rhythms of these substrates in the ocean ecosystem.

### Niche partitioning of purine and pyrimidine usage among heterotrophic bacterioplankton

We next sought to understand the ecological consequences of purine and pyrimidine excretion by *Prochlorococcus*. To this end, we searched for purine and pyrimidine usage genes in 305 partial and complete genomes of SAR11, SAR86 and SAR116, each among the most abundant of *Prochlorococcus*’ sympatric heterotrophs (16, 31–33). In SAR11, the occurrence of purine usage genes reaches frequencies similar to the average genome completeness of ∼72%, suggesting purine usage is a universal or nearly universal trait in this genus, whereas pyrimidine usage genes occur at frequencies of only ∼3% (Fig. 3, Data S1). In contrast, in SAR86, thymidine usage genes occur at frequencies similar to the average genome completeness of ∼71%, suggesting thymidine usage is a universal or nearly universal trait in the genus, whereas purine usage genes are absent (Fig. 3, Data S1). Finally, in SAR116, purine usage genes occur at frequencies of ∼49%, whereas pyrimidine genes occur at frequencies of ∼43% (Fig. 3, Data S1), both below the average genome completeness of ∼75%. These observations suggest that resource niche partitioning within the oceanic bacterioplankton community has given rise to purine and pyrimidine specialists, as well as generalists that use both classes of compounds, in turn driving differentiation of cross-feeding with *Prochlorococcus*.

**Fig. 3.**
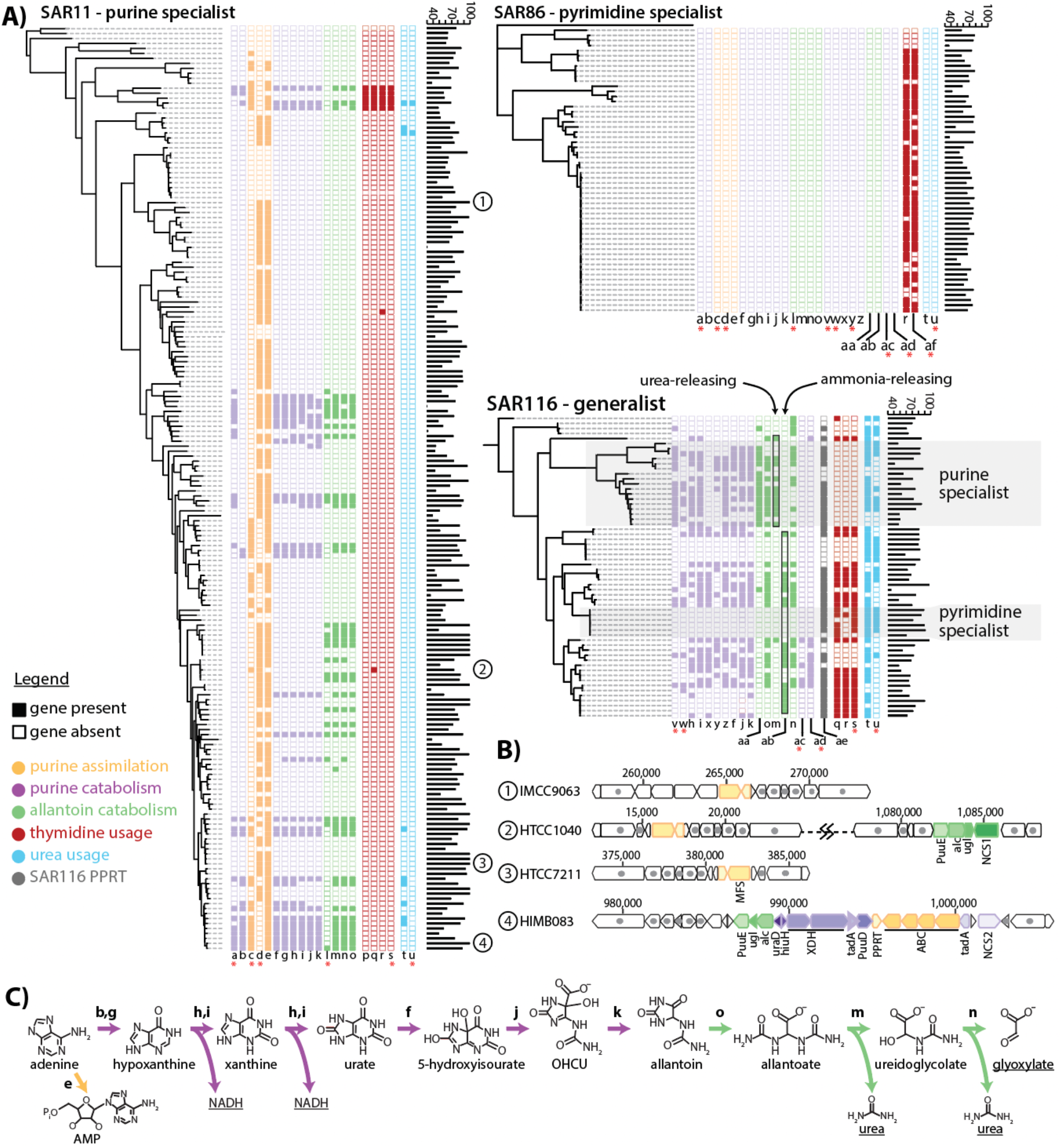
Niche partitioning of purine and thymidine usage in oceanic bacterioplankton. **A)** Phylometabolic diagrams show presence/absence of purine and pyrimidine usage genes in the abundant heterotrophic bacterioplankton groups SAR11, SAR86 and SAR116. Transporter genes are indicated with pink asterisks. Gene colors indicate different pathway functions as shown in the inset legend. Genome completeness statistics (%) are shown as bar graphs next to gene profiles. **B)** Representative genome profiles of SAR11 sub-groups specializing in either purine assimilation (IMCC9063 and HTCC7211), purine assimilation and allantoin catabolism (HTCC1040), or full purine catabolism (HIMB083). **C)** Purine catabolism pathway in SAR11, with major degradation products underlined: energy in the form of NADH, carbon in the form of glyoxylate and nitrogen in the form of urea. Genes: a = purine NCS2 permease, b = purine deaminase 2 (tadA), c = purine ABC transporter permease, d = purine MFS permease, e = purine phosphoribosyltransferase (PPRT), f = urate oxidase (PuuD), g = purine deaminase 1 (tadA), h = Xanthine dehydrogenase Mo-subunit (XDH_Mo), i = xanthine dehydrogenase FAD-subunit (XDH_FAD), j = 5-hydroxyisourate lyase (hiuH), k = 2-oxo-4-hydroxy-4-carboxy-5-ureidoimidazoline decarboxylase (OHCU decarboxylase, uraD), l = allantoin NCS1 permease, m = allantoicase (alc), n = ureidoglycolate lyase (ugl), o = allantoinase (PuuE), p = uracil phosphoribosyltransferase, q = cytidine deaminase, r = thymidine kinase, s = pyrimidine ABC transporter, t = urease, u = urea ABC transporter, v = purine ABC transporter, w = purine ABC transporter, x = XDH accessory protein, y = purine NCS2 permease, z = purine deaminase, aa = allantoin racemase, ab = ureidoglycine aminohydrolase, ac = purine TRAP transporter, ad = DMT transporter, ae = purine/pyrimidine phosphoribosyltransferase, af = CNT transporter.

Niche partitioning of purine and pyrimidine cross-feeding occurs across multiple taxonomic scales. For example, while at the genus-level SAR116 is a purine and pyrimidine generalist, it also contains subgroups that have genes for only purine or pyrimidine usage, but not both (Fig. 3, Fig. S2). In addition, different parts of the SAR116 tree contain variations in the pathway of purine catabolism that are characterized by differences in the presence or absence of allantoin racemase and the production of either ammonia or urea in the final steps of the pathway (Fig. 3, Fig. S3), as well as an association with primary, ATP-consuming transporters or with secondary, ion-gradient-driven transporters (Fig. 3).

Similarly, different SAR11 sub-groups have gene profiles suggesting they use purines for different purposes. For example, most genes making up a large operon for purine catabolism (34) occur in ∼18-22% of genomes, but an uptake transporter and a purine phosphoribosyltransferase that together perform the initial assimilatory portion of the pathway occur at frequencies similar to the average genome completeness of ∼72% (Fig. 3, Data S1), suggesting they are universal or nearly universal. This suggests that some SAR11 cells break purines down to their constituent components, while others use them intact. Further, an allantoin transporter occurs in ∼12% of genomes (Data S1), including in at least 10 genomes that lack the full purine catabolism operon but retain genes involved in allantoin catabolism (Fig. 3), leading to an overall higher frequency of allantoin catabolism genes (∼22%) than upstream purine catabolism genes (∼18%) in SAR11 (Data S1). Allantoin is the product of the first sequence of reactions of purine catabolism in which one of the rings is cleaved and energy is released, while its further degradation releases carbon and nitrogen (Fig. 4). It has been shown that some benthic algae can use allantoin (35) and that sponges produce it (36), but in general this metabolite has been little studied in marine systems. Mammals have been observed to excrete allantoin (37), while in plants, allantoin is transported across organelles during purine degradation (38) and is thought to be important for combating oxidative stress (39). Given the global abundance of SAR11 (31,33), the occurrence of an allantoin-using subgroup (Fig. 3) suggests that allantoin could have a broader role in marine ecosystems than currently recognized. Indeed, SAR11 clades often co-occur in the same environment (40), raising the possibility that in some environments SAR11 cells in search of energy incompletely degrade purines, releasing allantoin that is taken up and further degraded by surrounding allantoin-specialist SAR11 cells.

**Fig. 4.**
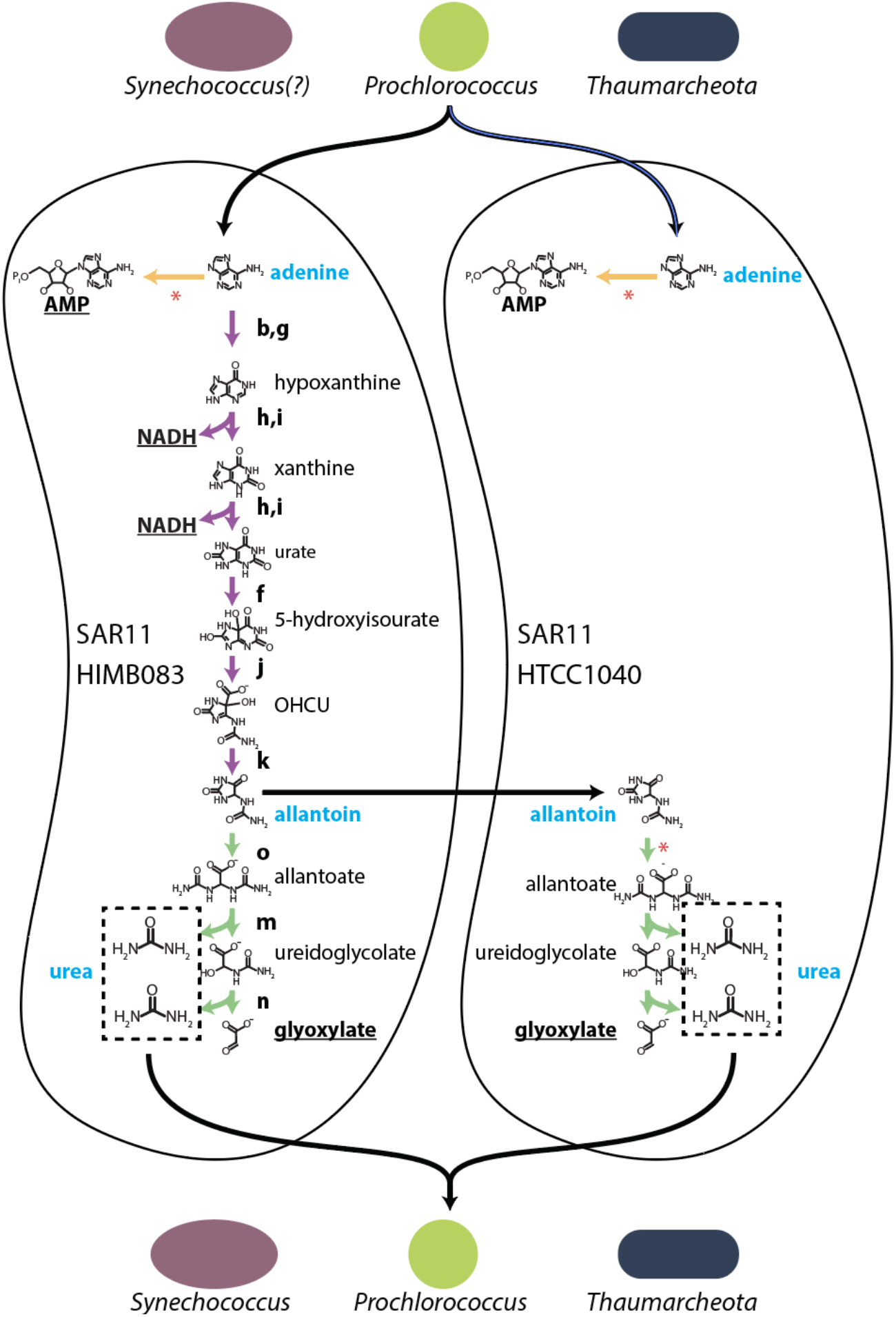
Putative purine:urea-mediated cryptic nitrogen cycles among abundant ocean microbes. *Prochlorococcus* and *Thaumarcheota* (and potentially *Synechococcus*, see text) supply purines that are used by SAR11. Different SAR11 cell types, labeled by representative strains (Fig. 3), possess variants of purine catabolism as shown using sequences of reactions that perform different purine usage functions. Underlined metabolites (AMP, NADH, glyoxylate) are the main products of purine assimilation/breakdown inferred to be used by the shown SAR11 cell types, while blue metabolites (adenine, allantoin, urea) are inferred to be involved in cross-feeding interactions. Many SAR11 genomes, including the examples shown, lack the urease genes required to use urea, suggesting it is released and available to surrounding cells of other groups, including *Prochlorococcus, Thaumarcheota* and *Synechococcus*. Some SAR11 genomes lack purine catabolism genes but contain genes for using allantoin, which is inferred to be released by other SAR11 cells that incompletely catabolize purines. Colors of purine usage functions are as defined in Fig. 3: purine assimilation genes in yellow, purine catabolism genes in purple and allantoin catabolism genes in green, while enzymes mediating different reactions are labeled with letters, also as defined in Fig. 3. Reactions marked with a red asterisk represent enzymes that in subsets of SAR11 genomes are collocated with transporter genes (Fig. 3). Black arrows represent inferred cross-feeding pathways. OHCU = 2-oxo-4-hydroxy-4-carboxy-5-ureidoimidazoline.

In contrast to the diversity in pyrimidine/purine usage functions observed in SAR116 and SAR11, nearly all SAR86 genomes contain just two, often-collocated thymidine usage genes: thymidine kinase and a thymidine transporter (Fig. 3, Fig. S2). SAR116 genomes, as well as some deep-branching SAR11 genomes, contain other genes for interconverting among pyrimidines that are collocated with thymidine kinase and the transporter, including uridine ribohydrolase, uracil phosphoribosyltransferase and cytidine deaminase, but these are absent in SAR86 (Fig. 3, Fig. S2). This suggests that while SAR116 may have a more general pyrimidine usage strategy, SAR86 focuses specifically on thymidine assimilation.

### Niche partitioning of purine usage strategies in SAR11

Since SAR11 is the most abundant of *Prochlorococcus*’ sympatric heterotrophs (31,33), we decided to examine its diversity of purine usage strategies more closely. As noted, purine catabolism in SAR11 leads to three products: energy in the form of NADH, carbon in the form of glyoxylate, and nitrogen in the forms of ammonia and urea (Fig. 3). However, the bulk of nitrogen released during purine catabolism is released as urea, whereas assimilation of nitrogen into biomolecules universally starts from ammonia. To use nitrogen in urea, cells across the tree of life must first break it down to ammonia using urease. We searched SAR11 genomes for urease genes, and found they occur at a much lower frequency (∼5%) than purine catabolism genes (∼18-22%) (Fig. 3, Data S1). This suggests additional niche partitioning of purine catabolism within SAR11, with some cells using it to obtain nitrogen, while others use it primarily to obtain energy and/or carbon, releasing urea back to the environment. Genomes in our sample that contain allantoin catabolism genes but lack the rest of the purine catabolism operon also lack urease (Fig. 3), similarly suggesting that these cells use allantoin as a carbon source and release urea back to the environment.

These observations suggest the existence of purine:urea-mediated cryptic nitrogen cycles involving many abundant microbial groups (Fig. 4). That is, in addition to exuding purines (21) (Fig. 1), *Prochlorococcus* is also a major urea user (41,42). This creates a putative cycle in which purines transfer energy and carbon from *Prochlorococcus* to SAR11 and urea transfers nitrogen back to *Prochlorococcus*. Incomplete purine catabolism by some SAR11 cells leading to potential transfer of allantoin to other surrounding SAR11 cells (Fig. 3 and surrounding discussion) may add an additional step to this cycle in some environments (Fig. 4). Similarly, marine *Synechococcus* also uses urea (41,42), and a marine strain of *Synechococcus elongatus* (which branches outside of marine picocyanobacteria) was found to exude thymidine (43), suggesting the deoxyribonucleotide recycling pathway we identified in *Prochlorococcus* (Fig. 2) could have a broader distribution across other cyanobacteria. Further, ammonia-oxidizing archaea (Thaumarcheota), which are highly abundant autotrophic (CO_2_-fixing) microbes lying at the base of the food web below the euphotic zone (44), also exude thymidine and purines (45) and use urea (46). Finally, SAR11 has a broad geographic range that extends across all these groups, from coastal to open waters and from the surface to the deep (40,47). Together these observations suggest that purine:urea-mediated cryptic nitrogen cycles (Fig. 4) may operate throughout much of the world’s oceans, linking many abundant groups, warranting their further study in both laboratory and field settings.

To further disentangle the forces shaping SAR11 purine usage functions and associated putative nitrogen cycles, we next examined the abundance of relevant genes in metagenomes collected from diverse oceans. We estimated the fraction of SAR11 cells with the capacity for different purine usage strategies in different environments by normalizing the abundance of purine usage genes to a set of single-copy core genes (Methods). We found that genes involved in purine assimilation are always abundant and have a relatively uniform distribution geographically, with depth, and as a function of nutrient concentrations (Fig. 5, Fig. S4-S9, Table S2), consistent with their near-universal presence in SAR11 genomes (Fig. 3).

**Figure 5.**
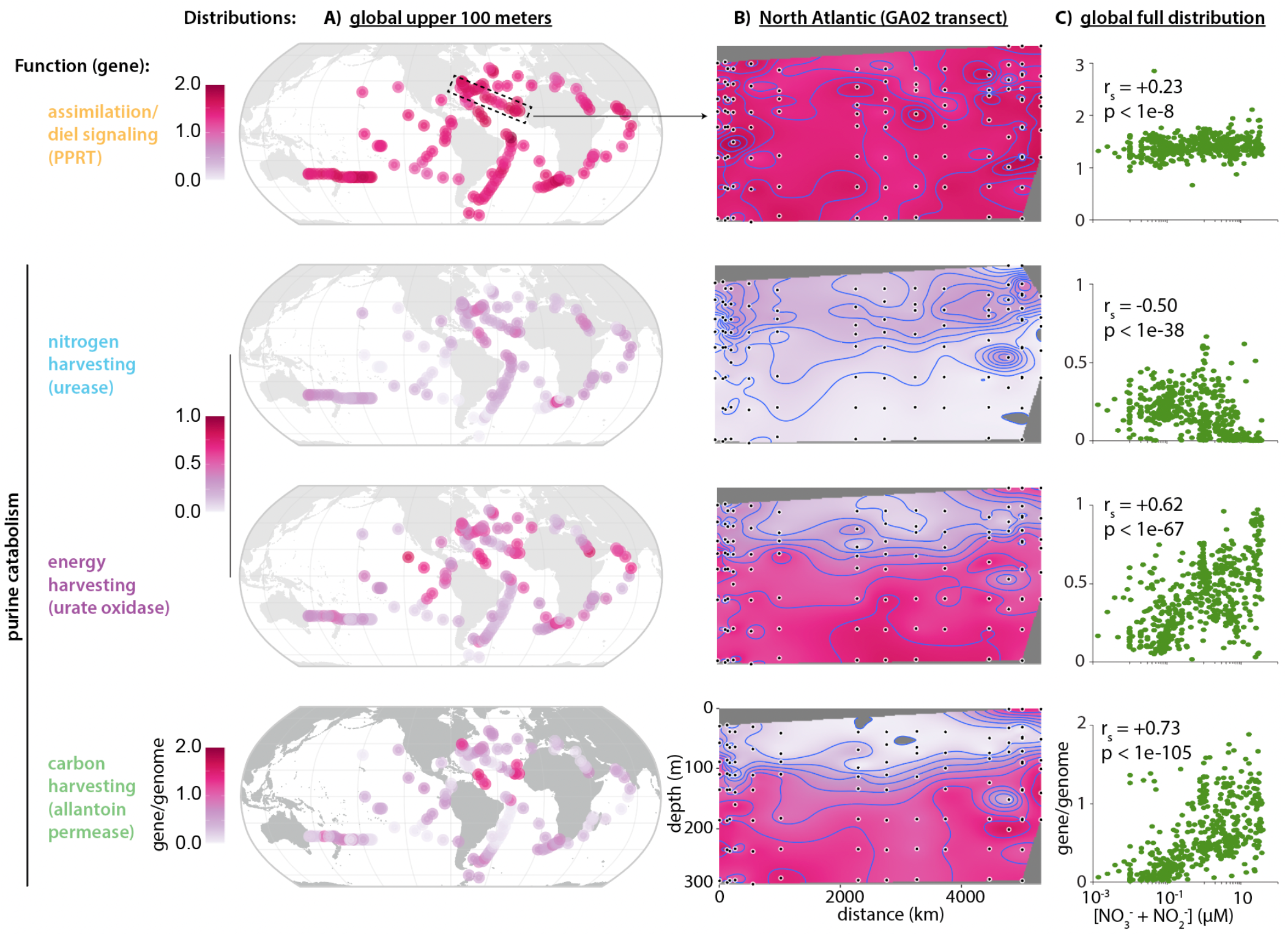
Distribution of SAR11 purine usage genes. Metagenomically-derived estimates for the gene/genome frequency of different purine usage genes in SAR11 are shown as a function of **A)** surface (i.e. upper 100 meters) biogeography, **B)** depth in the water column in the North Atlantic, and **C)** inorganic nitrogen concentration. Purine usage functions are color coded as defined in Fig. 3. For surface biogeography panels each data point represents the mean across samples within the upper 100 meters of the water column at a given station. Spearman rank correlation coefficients (r_S_) of the correlation between gene/genome frequency and nitrogen concentration are shown as insets in panels on the right. PPRT = purine phosphoribosyltransferase.

Genes mediating other purine usage functions in SAR11 have distinct, non-uniform distributions: genes of the carbon– and energy-harvesting portions of the pathway increase in frequency in environments where nutrients are elevated, including in surface regions in the Arabian Sea, the upwelling regions of the Eastern Equatorial Pacific and the Benguela system off the tip of South Africa, and the Amazon River plume off the northern coast of South America (Fig. 5, Fig. S4-S9). Genes involved in harvesting nitrogen from purines follow the opposite pattern, reaching their highest abundances in environments where nutrients are depleted (Fig. 5, Fig. S4-S9). While energy and carbon harvesting can proceed without the downstream process of nitrogen harvesting, nitrogen harvesting can only proceed if the full pathway is present (Fig. 4). Hence, the similar abundance of genes for both functions in the nutrient-poor open ocean – where the abundances of genes for carbon/energy-harvesting reach their minimum – suggests that in those environments SAR11 primarily uses purine catabolism to obtain nitrogen.

Further insight into the forces underlying niche partitioning come from the depth distribution of purine catabolism functions in SAR11, as seen most clearly in the North Atlantic (Fig. 5, Fig. S5-S9). That is, both energy– and carbon-harvesting genes are most abundant at depth, where inorganic nutrient concentrations increase but the energy supply to heterotrophs from primary producers decreases. In contrast, nitrogen-harvesting genes are most abundant near the surface where energy is more available, but nutrients are depleted. Taken together these observations suggest that niche partitioning of SAR11 purine usage functions occurs along a central axis defined by the relative availability of nitrogen and energy. This is consistent with the different transporters we identified in SAR11 as being associated with the different functions of purine catabolism in this group: nitrogen-harvesting genes are associated with an ABC transporter, a high-affinity but energetically costly system requiring ATP hydrolysis (48), while both energy– and carbon-harvesting genes are associated with more energetically efficient nucleotide:cation symporters (49) (Fig. S4-S9, Table S2).

Genes for the allantoin transporter characteristic of the allantoin-using SAR11 subgroup (Figs 3,4) follow a broadly similar trend as genes involved in energy-harvesting but are nearly absent at the surface where energy-harvesting genes remain present (Fig. 5, Fig. S5-S9). The frequency of carbon-harvesting genes also has a stronger positive correlation with nutrient concentrations than the frequency of energy-harvesting genes (Fig. 5, Fig. S4, Table S2). These observations are consistent with a decoupling of energy– and carbon-harvesting portions of the purine catabolism pathway in some SAR11 cells as the total flux of purine catabolism increases, thereby making allantoin available to other surrounding cells (Fig. 4). This is further consistent with the use of symporter systems to acquire purines and allantoin at depth, as it suggests that levels of both are higher in deeper waters than at the surface.

While phylogenomic and metagenomic analyses identify patterns in the genomic capacity of groups potentially involved in cross-feeding, they do not directly address the interaction between specific groups. In long-term co-cultures of SAR11 and *Prochlorococcus*, in turn, their growth rates match each other while their relative abundance stabilizes (33), both of which are consistent with SAR11 becoming dependent on *Prochlorococcus* exudates, but the pathways involved have not been unraveled. Indeed, metabolic processes in surface oceans play out over diel light:dark cycles, which can lead to temporal niche partitioning (50). We therefore reexamined existing in situ community gene expression data taken over several diel light:dark cycles in the North Pacific Ocean (51) in search of additional evidence regarding cross-feeding of purines between *Prochlorococcus* and SAR11. We found that expression of the SAR11 purine transporter that we infer is involved in purine assimilation (i.e. yellow genes in Figs. 3 & 4) closely follows in time expression of *Prochlorococcus* RNA polymerase genes, with a nighttime maximum a few hours after peak expression of *Prochlorococcus* DNA polymerase genes (Fig. 6, Data S2). Since we inferred that the transition from DNA to RNA synthesis is when Prochlorococcus releases purines (Fig. 2), these patterns are consistent with cross-feeding of purines between the two genera.

**Fig. 6.**
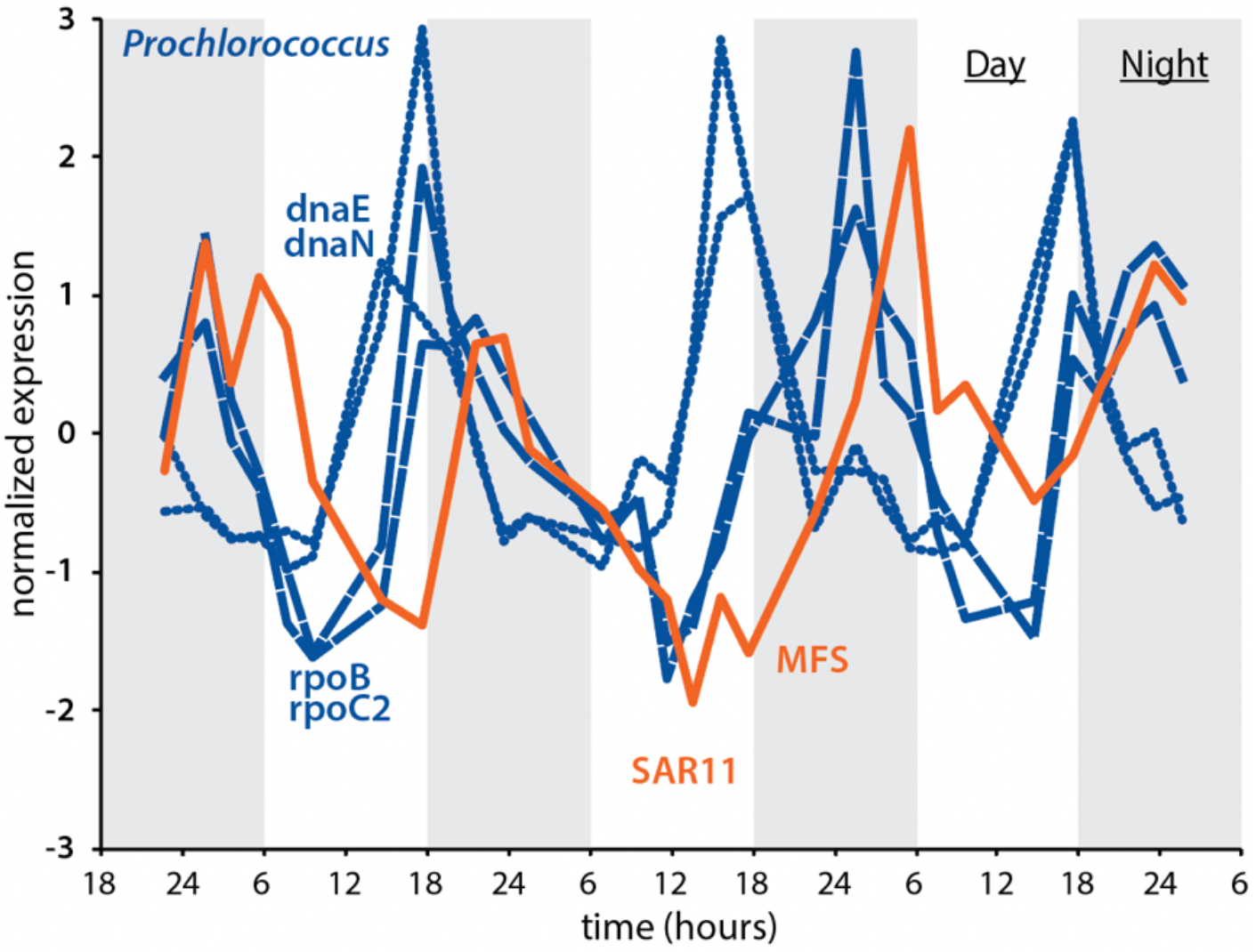
In situ expression of genes related to putative purine cross-feeding between *Prochlorococcus* and SAR11. Normalized in situ expression of DNA polymerase (dnaE, dnaN) and RNA polymerase (rpoB, rpoC2) in *Prochlorococcus*, and of a SAR11 purine MFS permease, in the North Pacific Ocean (data from Ref. 51). *Prochlorococcus* genes are in dark blue, while the SAR11 transporter gene is in orange. The transition from DNA to RNA polymerization in Prochlorococcus is inferred to be associated with the release of purines (Fig. 2), while the SAR11 purine MFS transporter is inferred to be involved in the intact assimilation of purines (Figs 3,4).

Transcripts of SAR11 purine catabolism genes have very low abundances in the data (Data S2), indicating the sequencing depth is insufficient to characterize patterns in their expression. We suspect this may in part reflect the observation that these genes reach frequencies <25% in metagenomes from surface waters of the North Pacific (Fig. 5, Fig. S5), where metatranscriptome samples were obtained (51). The low abundance of SAR11 purine catabolism genes and transcripts coinciding with a maximum in the frequency of SAR11 urease genes in surface waters (Fig. 5, Fig. S5-S9) further suggests that any flux of purine catabolism that does occur in SAR11 populations in that environment is primarily involved in nitrogen harvesting, resulting in a small return of urea from SAR11 to Prochlorococcus (Fig. 4). Indeed, expression of both urea transporter and urease genes in *Prochlorococcus* reach a maximum prior to the maximal expression of SAR11 purine assimilation genes (Fig. S10), suggesting it obtains urea primarily from sources other than SAR11. Isotopic labeling studies and more deeply sequenced metatranscriptome data from both surface waters and from deeper in the water column, as well as from other regions of the ocean with higher frequencies of purine catabolism genes in SAR11 genomes, will help constrain variations in the flux through different components of putative cross-feeding interactions (Fig. 4).

Our findings suggest potentially generalizable principles of metabolic evolution in microbial ecosystems. That is, niche partitioning describes how ecologically similar species may coexist in space and time by using their shared environment differently (52). In microbes this differentiation can be facilitated by acquiring traits via horizontal gene transfer, decoupling the evolutionary history of these traits from the parent species, suggesting that in some cases the genes themselves may be more appropriate units of study for understanding processes of ecological differentiation. For example, sub-populations of *Prochlorococcus* and SAR11 in different parts of the ocean are differentially enriched in genes for nutrient stress and nutrient assimilation due to local conditions (53–55), but individuals from these populations are interspersed with one another in phylogenies of universal core genes (53,55). Here, we find a similar pattern for genes involved in purine usage in SAR11, with their differentiation being clearer in terms of their distribution in the environment (Fig. 5) than in terms of their distribution across the SAR11 tree (Fig. 3). Together, these findings suggest that metabolic pathways – in this case involved in cross-feeding interactions among microbial species – can themselves in effect undergo a process of niche partitioning, driven by the biochemical and ecological tradeoffs between functions. This is consistent with the ‘It’s the song, not the singer’ framework of evolution, which proposes “casting metabolic and developmental interaction patterns, rather than the taxa responsible for them, as units of selection” (56).

While patterns of niche partitioning of SAR11 purine usage functions are clearest at the level of the genes themselves, these patterns are also still linked to the niche partitioning of taxonomically-defined clades, adding further support to our conclusions. For example, clades IC and IIB, which dominate in the mesopelagic (40,47) and upper mesopelagic (40,57), respectively, have higher frequencies of genes involved in harvesting energy (∼71% in clade IC, ∼47% in clade IIB) and carbon (∼57% in clade IC, ∼44% in clade IIB) from purines than other clades, while also having a high frequency of allantoin transporter genes (∼57% in clade IC, ∼50% in clade IIB), but lacking urease genes (Fig. S11, Table S3). This is consistent with our conclusions that the harvesting of energy and carbon from purines and the cross-feeding of allantoin become more prominent at depth, whereas nitrogen harvesting from purines is restricted to surface environments (Fig. 5 and surrounding discussion). Genomes of clade IA.1, which is abundant in cold surface waters that are typically richer in nutrients than warmer waters (58), in turn, have high frequencies of genes involved in allantoin transport (∼57%) and catabolism (∼71%), but lack genes involved in harvesting energy from purines. This is consistent with our conclusion that in some environments other SAR11 cells more focused on obtaining energy from purines make allantoin available to surrounding cells (Figs 3 and 4 and surrounding discussion). Further, genomes of clade IIIB, which represents a freshwater clade of SAR11 (59), lack all purine catabolism genes, consistent with purine cross-feeding being a feature specific to oceanic populations. Finally, clades IV, IIIA and IIA, all of which are abundant in surface waters in the Northwestern Atlantic from spring through early fall when waters are relatively more stratified (40), lack all genes for purine catabolism but retain genes for purine assimilation (Fig. S11, Table S3), consistent with our conclusion that this function is key in oligotrophic surface waters. Together these patterns suggest that differentiation of purine usage strategies played a role in the broader metabolic differentiation of SAR11 clades.

### Inferred role of purines in the synchronization of ocean-surface populations of SAR11 to daily rhythms in the supply of carbon from photosynthesis

Phylogenomic and metagenomic analyses of the functions of purine usage genes in SAR11 leave a key question unaddressed: what is the function of purine assimilation (yellow pathway in Figs. 3,4) in cells that lack the downstream purine catabolism pathway? In surface waters of the open ocean, where *Prochlorococcus* dominates, the frequency of purine catabolism genes in SAR11 genomes generally remains below ∼30-35% (Fig. 5, Fig. S5-S9), suggesting that assimilation of intact purines is the major form of purine usage in these populations (Fig. 6). To examine possible benefits of purine assimilation, we cultured SAR11 HTCC7211, a strain possessing assimilation genes but lacking purine catabolism genes (Fig. 3), in media amended with adenine at a range of concentrations. Since purines are nitrogen-rich and our metagenomic analyses suggest that SAR11 cells near the surface are often in search of nitrogen (Fig. 5), we performed these experiments in both nutrient-balanced (pyruvate:glycine:methionine = 1:1:0.2) and glycine-depleted (pyruvate:glycine:methionine = 50:1:10) media (Methods). We expected that by lowering the need for *de novo* purine synthesis, SAR11 cultures might receive a modest growth boost from the adenine additions, reflected either as an increased growth rate and/or a higher final culture density. However, we did not observe growth stimulation, but rather a decrease in growth rate at higher adenine concentrations (Fig. 7). The decrease in growth rate occurred at lower adenine concentrations in glycine-depleted than in nutrient-balanced cultures (Fig. 7).

**Figure 7.**
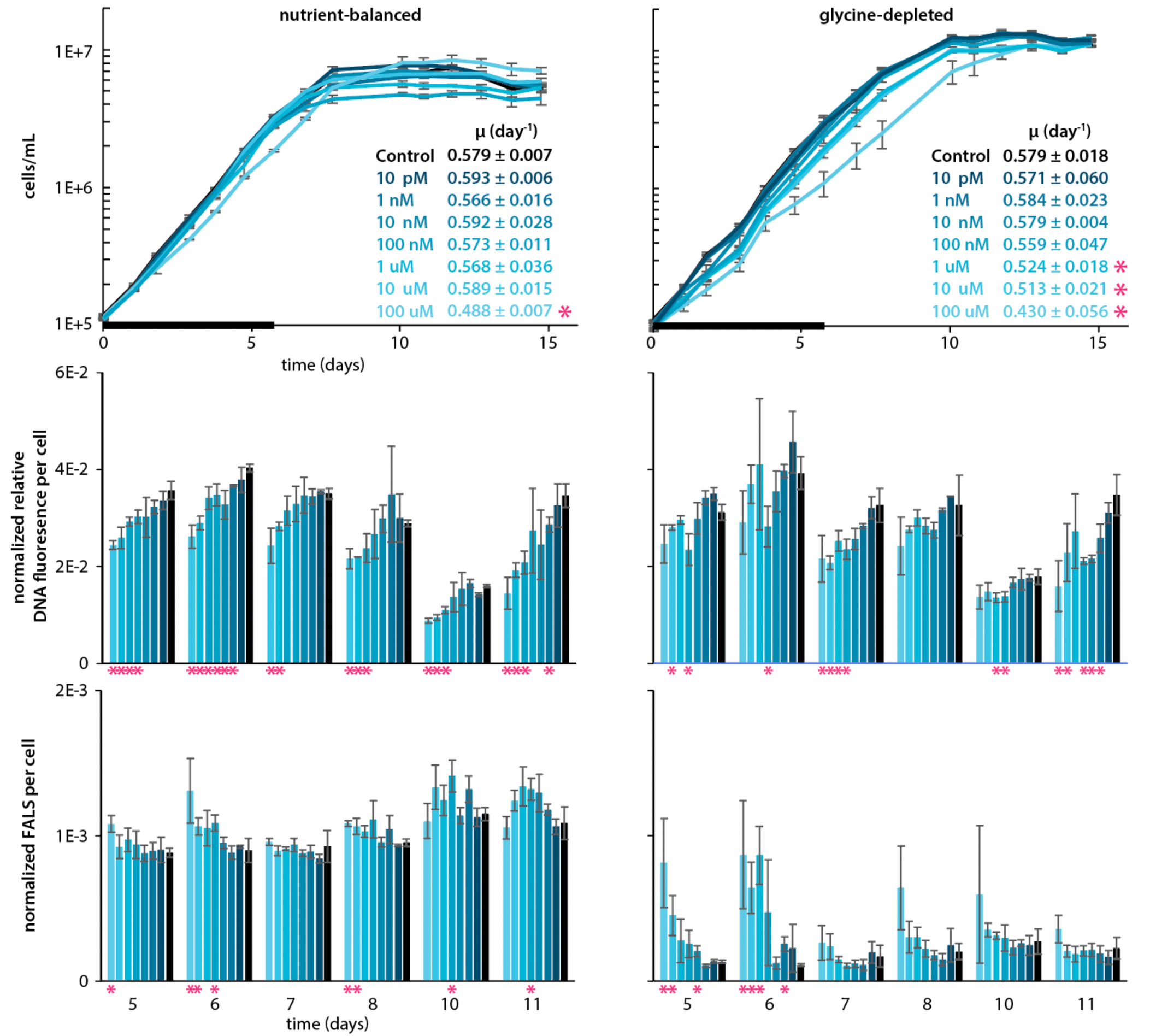
Growth and cell physiology of SAR11 HTCC7211 grown with and without adenine. Top panels: growth curves of SAR11 cells in nutrient-balanced ([pyruvate]:[glycine]:[methionine] = 1:1:0.2) and glycine-depleted ([pyruvate]:[glycine]:[methionine] = 50:1:10) media amended with different concentrations of adenine (shades of blue). Black bars at bottom reflect the time frame used to determine growth rates, which are shown in the insets. Middle and bottom panels: histograms of the averages of bead-normalized relative DNA fluorescence per cell and bead-normalized forward angle light scatter (FALS) per cell, respectively, at several time points between mid-exponential growth and stationary phase. For all panels, error bars reflect the standard deviation across three biological replicates, while values significantly different (p<0.05, 2-tailed Student’s t-test) between adenine-treated and unamended controls are highlighted with a pink asterisk.

Puzzled by these results, we wondered if adenine could provide an indirect benefit to cells lacking purine catabolism. A potentially relevant clue comes from *in situ* studies of genes expression, which indicate that in warm oligotrophic waters where *Prochlorococcus* dominates, SAR11 metabolism is synchronized to the diurnal light:dark cycle (51) despite not being a photoautotroph, and this coupling dissipates in nutrient-enhanced upwelling regions with higher nutrient levels and fewer *Prochlorococcus* (60). Further, negative feedbacks are often central in the synchronization of biological systems (61–63), and in eukaryotic systems purine and pyrimidine additions have been used to synchronize cell cultures (64–66). These purine and pyrimidine “block” techniques rely on the fact that the specificity of ribonucleotide reductase, which generates all dNTP precursors to DNA (Fig. 2), is controlled through allosteric binding of dNTP molecules (67). Consequently, it has been shown in eukaryotic systems that imbalances in intracellular dNTP pools due to pyrimidine/purine amendments can drive depletion in one or more dNTPs, thereby slowing down or fully inhibiting DNA synthesis and genome replication (65). Subsequent removal of the pyrimidine/purine block amendments by washing cultures results in cells undergoing genome replication and cell division in synchrony (64,65). If similar or related mechanisms play out in bacteria, this raises the possibility that inferred daily pulses of purines from *Prochlorococcus* (Fig. 2 and surrounding discussion) could act as a signal that helps SAR11 synchronize its metabolism to the daily rhythms of primary production.

To explore this hypothesis, we first examined flow cytometric data from SAR11 cultures amended with adenine to better understand how adenine inhibits SAR11 growth. Cells in SAR11 cultures amended with adenine have a lower average DNA content, and DNA content decreases as adenine concentrations increase (Fig. 7, Fig. S12). Further, cells from adenine-amended cultures are also larger at the higher concentrations of adenine that inhibit growth (Fig. 7). This increase in size is particularly noticeable in glycine-depleted cultures where growth inhibition sets in at lower adenine concentrations (Fig. 7). Together these results suggest that adenine additions inhibit DNA synthesis in SAR11, which, as the inhibition strengthens, leads to inhibition of cell division. This is similar to how purine and pyrimidine “blocks” function in eukaryotic systems (64–66).

In eukaryotic systems, purine and pyrimidine blocks in which DNA synthesis has been inhibited lead to cells being poised for division (64,65). Hence, we next examined whether SAR11 cells whose DNA synthesis has been inhibited by adenine are poised to divide. Since plating-based techniques or microscopic tracking of division in single cells are currently intractable in SAR11, we instead opted for a population-level approach to testing this possibility. That is, we grew SAR11 cultures in media with and without adenine, washed cells in mid-exponential growth, and compared their growth after resuspending in adenine-free media. Control cultures grown without adenine often experienced significant growth retardation after washing, with >60% of replicates experiencing major lags and/or never fully recovering (Fig. 8), suggesting that the ultra-centrifugation-based washing protocol induces stress in SAR11 cells. The reasons for the significant divergence of growth trajectories after washing among replicates are not clear, but it was a highly repeatable hallmark of cultures not exposed to adenine. In contrast, cultures treated with adenine display fewer signs of stress after washing, with <20% of replicates experiencing lags or failing to resume growth under most conditions tested (Fig. 8). In nutrient-balanced cultures the benefit of adenine additions is clearest at higher adenine concentrations, while in glycine-depleted cultures the benefit of adenine additions was clear across the full range of concentrations we examined (Fig. 8). Higher adenine concentrations, as well as adenine amendments in glycine-depleted relative to nutrient-balanced cultures, are both associated with greater inhibition of DNA synthesis and cell division (Fig. 7). More consistent reignition of growth after washing of SAR11 cultures experiencing greater inhibition of DNA synthesis is consistent with the inhibition poising cells for replication. However, we cannot fully rule out that other possible mechanisms underpin the indirect benefit of adenine in our experiments, particularly as it has other metabolic roles, including as the backbone for the central energy currency ATP. Indeed, while DNA synthesis is partially inhibited in SAR11 under a wide range of adenine concentrations, as evidenced from a decrease in DNA/cell, growth is not impacted at most concentrations we tested (Fig. 6). This suggests that more consistent reignition of growth after washing in those cases (Fig. 7) could be due to energetic priming of metabolism rather than poising of cells for division.

**Figure 8.**
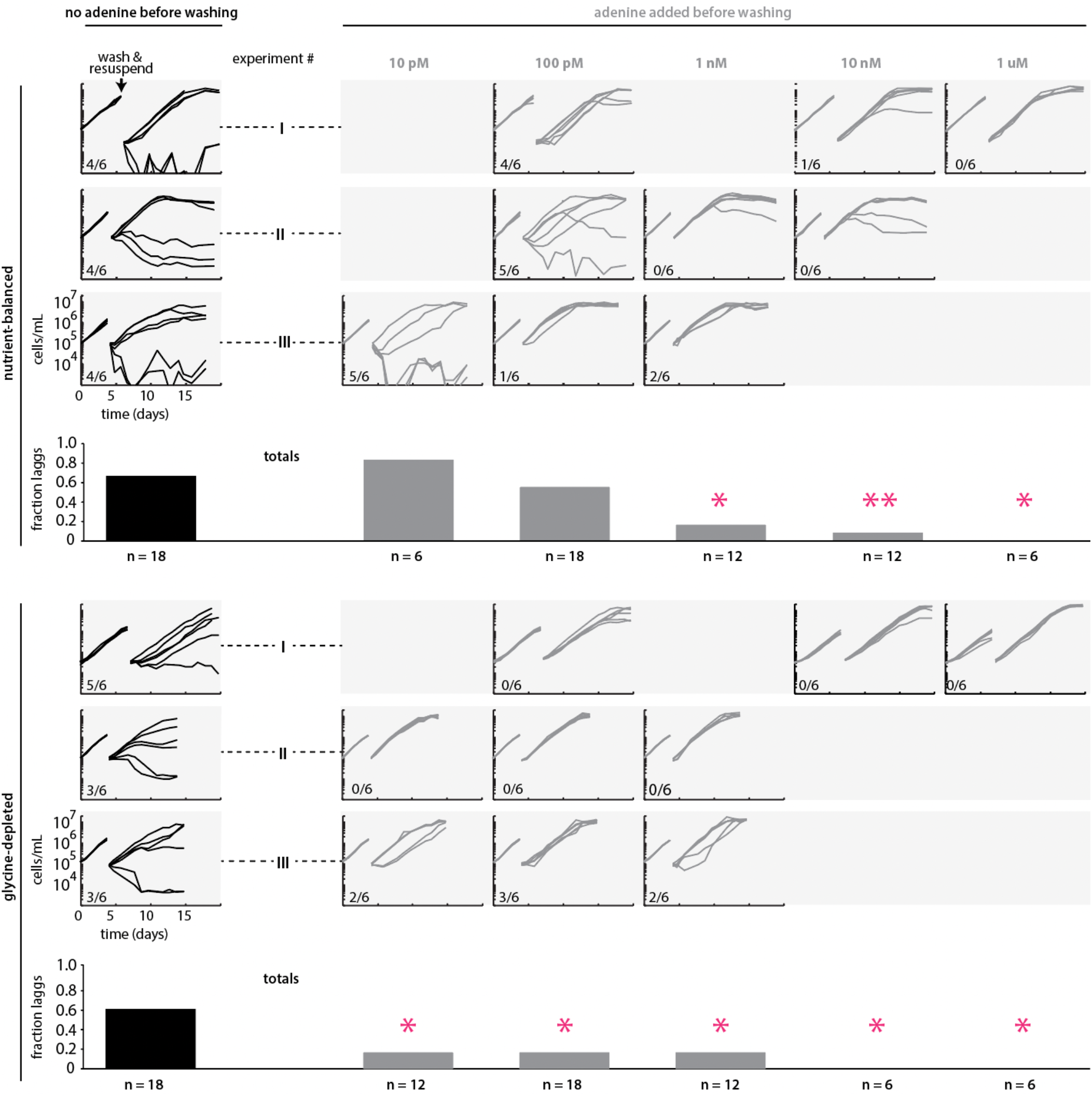
Post-washing response of SAR11 HTCC7211 previously grown with and without adenine. Growth curves of SAR11 cultures grown with and without adenine, before and after washing and resuspension in adenine-free media. Three identical experiments were performed in both nutrient-balanced and glycine-depleted cultures, which are labeled using roman numerals. Each experiment included cultures grown with adenine at three different concentrations as well as an adenine-free control, with 6 biological replicates for each condition. Empty entries at given concentrations in different rows represent adenine concentrations not included in different experiments. Fractions of replicates experiencing a lag of 2 or more days after washing at a given adenine concentration and culture condition are shown as inset within each experimental panel, and as fractions of totals across experiments at bottom. Statistically significant differences between adenine amended and adenine-free cultures according to a 2-tailed Fisher’s exact test are shown as single pink asterisks for p<0.05 and double pink asterisks for p<0.01.

From these observations, a conceptual model emerges in which SAR11 populations in surface waters of the open ocean use purines from *Prochlorococcus* as a signal to synchronize to daily rhythms in the supply of energy and carbon from photosynthesis (Fig. 9). While it is not clear if negative inhibition from adenine is absolutely required to achieve synchronization, we summarize the evidence in favor of the hypothesis that it plays this role: 1) absent an indirect benefit, we lack an explanation for why natural selection maintained purine assimilation genes in SAR11 when other purine catabolism genes were lost (Fig. 3), 2) biochemical oscillators generally rely on some form of negative feedback to achieve sustained synchronicity due to its role in returning the system to a given baseline state (61–63), and we experimentally detect inhibition of DNA synthesis by exogenous purines in SAR11 (Fig. 7), 3) we observe a correlation between the strength of inhibition of DNA synthesis and the likelihood of growth reignition in SAR11 cultures experiencing stress (Fig. 8), suggesting the inhibition of DNA synthesis propagates through cellular physiology, priming metabolism and/or poising cells for division, 4) we infer from metabolic reconstructions (Figs. 1,2) that *Prochlorococcus* releases nightly pulses of purines, providing a potential ecological timing mechanism to SAR11, 5) in situ community gene expression data (51) shows that expression of the inferred SAR11 purine assimilation transporter closely follows that of *Prochlorococcus* metabolism genes inferred to be associated with purine release (Fig. 6), 6) apparently similar mechanisms based on inhibition of DNA synthesis by exogenous pyrimidines and purines have been used to synchronize eukaryotic cell cultures (64–66). Together, this leads us to propose a model (Fig. 9) in which the daily alternation of positive and negative cross-feeding signals increases the effective metabolic coupling of SAR11 to the daily solar energy supply, thereby providing the selective pressure to maintain purine assimilation genes when other catabolism genes were lost.

**Fig. 9.**
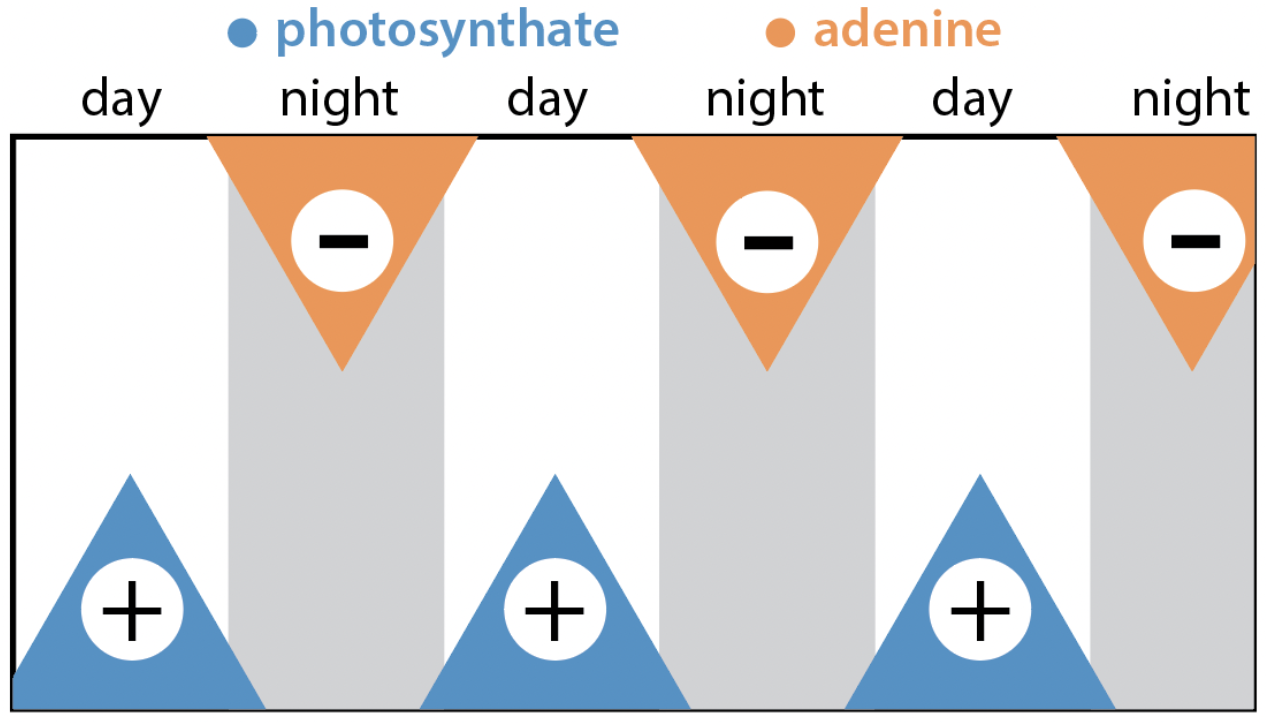
Hypothesized role of purines in driving synchronization of SAR11 metabolism to the diel light:dark cycle. Organic carbon produced by phytoplankton during the day (photosynthate, blue) stimulates SAR11 metabolism, while purines (orange) exuded by *Prochlorococcus* at night after genome replication (Fig. 2) inhibit SAR11 metabolism (Fig. 6). Daily alternation of these positive and negative signals over long timescales drives synchronization of metabolism to the 24-hour period as observed in *in situ* gene expression data (51).

The degree to which diel light:dark synchronization extends from metabolism to cell division in wild populations of SAR11 is an open question. First, as pointed out above, at lower concentrations likely to be more ecologically relevant, adenine inhibits DNA synthesis but not growth of SAR11 (Fig. 7), and still promotes greater reignition of growth after washing (Fig. 8), suggesting metabolic effects not directly linked to cell division could be at play. Further, metagenomics-based computational approaches that detect known diel replication patterns in *Prochlorococcus* do not detect them in SAR11 (67), suggesting that if diel replication does occur in the latter, it is less pronounced. Lastly, in co-cultures maintained in balanced growth over many transfers, the growth rates of *Prochlorococcus* and SAR11 match each other and their relative abundances remain stable (33), suggesting a coupling of their metabolisms, but patterns of division have not been characterized for growth under diel light:dark conditions. Further studies of daily cellular replication patterns in SAR11, both in the field and in laboratory co-cultures with *Prochlorococcus*, are key areas of future research that will deepen our understanding of mechanisms of synchronization within oceanic microbial ecosystems.

In the context of our proposed framework (Fig. 9), it is worth reconsidering SAR86’s genomic profile for using thymidine, which our metabolic reconstructions suggest is also released by *Prochlorococcus* in nightly pulses (Fig. 2). That is, SAR86 has a genomic profile that suggests it focuses specifically on assimilating – and not interconverting, or breaking down – thymidine (Fig. 3), and in that sense is like SAR11 populations that possess purine assimilation genes but lack purine catabolism genes. Further, as in SAR11, *in situ* gene expression studies indicate that SAR86 metabolism is synchronized to the diurnal light:dark cycle in oligotrophic waters (51), but not in regions of upwelling where *Prochlorococcus* is less abundant (60). This raises the possibility that thymidine could play a similar metabolic role in SAR86 that adenine plays in SAR11. Our findings highlight the potential value of studies of the dynamics of dissolved purines and pyrimidines in wild microbial communities and in microbial cultures, as well as of the coupled dynamics of *Prochlorococcus* and sympatric heterotrophs.

## Materials and Methods

### Determining the scale of pyrimidine and purine production and exudation by Prochlorococcus

Data on the intra– and extracellular abundance of metabolites in *Prochlorococcus* were obtained in Ref. (21) (also available at MetaboLights (https://www.ebi.ac.uk/metabolights/MTBLS567). Briefly, in our previous study (21), spent media was obtained in mid-exponential growth from nutrient replete batch cultures of *Prochlorococcus* strains MIT9301, MIT0801 and MIT9313 grown at multiple light levels, as well as from semi-continuous P-limited cultures of *Prochlorococcus* MIT9301. Intracellular metabolites were directly extracted from cells using organic solvents and water. Metabolites from the spent media were first extracted using solid phase extraction. Extracts were then analyzed using UPLC-MS/MS to determine concentrations. For extracellular metabolites, concentrations in the extracts were corrected for the extraction efficiency using the formula: [*m*]_media_ = 100 x [*m*]_extract_ /EF%, where [*m*]_media_ is the concentration of metabolite *m* in the spent media, [*m*]_extract_ is the measured concentration in the extract, and EF% (range = 0 – 100) is the efficiency with which the solid phase extraction resin extracts metabolite *m* from seawater (68). This correction is only performed for metabolites with EF% > 1 due to associated uncertainties in precisely determining extraction efficiencies when they are low (21). While thymidine and methylthioadenosine have an EF%>1, adenine and guanine have an EF%<1, and so their concentrations are left uncorrected.

Next, extracellular concentrations of purines and thymidine were normalized to the total amount of these compounds incorporated into DNA over the course of the experiment using experimentally determined cell counts and known GC% of the genomes for each strain, and assuming one chromosome per *Prochlorococcus* cell (29). Thymidine, adenine and methylthioadenosine were normalized to the AT% of genomes, while guanine was normalized to the GC% of genomes.

For intracellular purines and pyrimidines, concentrations were measured in units of ng/mL in the cell extract and converted to fg/cell using the extract volume and the total number of cells captured on the filter. Total cell numbers are obtained by multiplying culture volume and culture density as determined by flow cytometry. Concentrations in units of fg/cell are then divided by the molar mass of metabolites to obtain concentrations in units of moles/cell. Finally, concentrations in units of moles/cell are divided by cellular volume to obtain molar concentrations. Cellular volumes for *Prochlorococcus* were determined as follows. The volumes of *Prochlorococcus* strains MIT9313 (0.44 µm^3^) and MED4 (0.2 µm^3^) were measured in Ref. (70), while the cellular dry masses of strains MIT9301 (60 femtograms), MED4 (66 femtograms), NATL2A (91 femtograms) and MIT9313 (158 femtograms) were measured in Ref. (71). The volume of MIT9301 was estimated from that of MED4 by scaling by the relative dry weight masses of the two strains (i.e. 60 fg/cell for MIT9301 and 66 fg/cell for MED4, giving a ratio of 60/66), leading to 0.18 µm^3^ for MIT9301. While there are no volume or mass measurements of MIT0801 it belongs to the same “low light adapted I” (LLI) ecotype as NATL2A and so we estimated its volume from that of MED4 by scaling by the relative dry weight masses of NATL2A and MED4 (i.e. 91 fg/cell for NATL2A and 66 fg/cell for MED4, giving a ratio of 91/66), leading to 0.28 µm^3^ for MIT0801.

### Reconstructing putative deoxyribonucleotide recycling pathway

To identify possible pathways involved in the production and/or consumption of thymidine, adenine and guanine, we searched the annotated genomes of all *Prochlorococcus* strains available on the KEGG database (72) for all possible genes involved in the metabolism of these compounds. All strains used in this study (MIT9301, MIT0801 and MIT9313) are included on KEGG. To place findings in a broader context we also searched all *Prochlorococcus* genomes for all genes involved in the metabolism of nucleobases, nucleosides, deoxynucleosides, ribonucleotide monophosphates and deoxyribonucleotide monophosphates, all intermediates of a larger nucleotide metabolic network that thymidine, adenine and guanine could be a part of. To account for the possibility of gene misannotations or incorrect functional assignments in some *Prochlorococcus* strains, homologs identified in some *Prochlorococcus* strains were used to query all *Prochlorococcus* genomes using the BLASTp algorithm within the KEGG BLAST feature, using an e-value cutoff of E < 1e-25. We included in our search Fatty Acid Metabolism-Immunity Nexus (FAMIN), a recently characterized enzyme that is highly multi-functional and catalyzes multiple reactions within nucleotide metabolism (73). Finally, to identify potentially relevant genes that are present but not functionally assigned in any *Prochlorococcus* genome, we also used the BLASTp algorithm within KEGG to search sequences of additional purine/pyrimidine metabolism genes from *E. coli* (which has an extensive network of nucleotide metabolism) against *Prochlorococcus* genomes using a permissive E-value cutoff of E < 1e-10, but this did not lead to identification of any further genes. Altogether we identified 9 genes, 5 of which are universal in all strains used in this study (Table S4). Of these 5 genes, only 3 (SurE, MTAP, apt) could be assembled into a putative pathway that involves purines and pyrimidines exuded by *Prochlorococcus*. Data on the temporal transcriptional profiles of all 9 genes were obtained from a study in which the growth of *Prochlorococcus* MED4 was synchronized to diurnal L:D conditions (29). To evaluate transcriptional dynamics of these genes relative to DNA and RNA synthesis, the transcriptional profile of each gene was plotted alongside the transcriptional profile of DNA polymerase and RNA polymerase (Fig. S1). Pearson correlation coefficients of the expression of each gene relative to DNA and RNA polymerase genes were calculated in Excel (Table S5).

### Phylometabolic analysis of purine and pyrimidine strategies in heterotrophs

Genomes, curated genome phylogenies and associated metadata for SAR11, SAR86 and SAR116 were obtained from the MARMICRODB database (74). The MARMICRODB database includes genomes from a variety of sources, including isolate genomes, single-cell amplified genomes (SAGs) and metagenome assembled genomes (MAGs), but genomes with %completeness – 5x %contamination < 30 were excluded in the final dataset, resulting in 186 SAR11 genomes, 57 SAR86 genomes and 59 SAR116 genomes. While exploring these genomes we noticed many contained operons of purine and pyrimidine usage genes (Fig. 3, Fig. S2). Homologs from those operons were used for local searches using the BLASTp algorithm obtained from NCBI (75), using an E-value cutoff of E < 1e-25. However, since some homologs with other functions occasionally fall below this cutoff and our genome set also included MAGs and SAGs that contain partial genes that can have higher E-values, final counting of genes relied on a combination of sequence similarity, E-values and genomic context (i.e. whether genes occurred in operons). For example, many SAR11 and SAR116 genomes that contain a complete purine catabolism operon contain a second homolog also annotated as 5-hydroxyisourate lyase, but which occurs in a different genomic location and has low sequence similarity to the homolog occurring in the operon. Since this gene further occurs in many genomes that do not contain any other purine catabolism genes, we conclude it has a different function and leave it out of our final tally. All purine and pyrimidine usage genes thus identified are listed, along with their gene/genome frequencies, in Data S1. Finally, presence/absence data of genes for individual genomes was mapped onto the leaves of the SAR11, SAR86 and SAR116 genome phylogenies to generate phylometabolic trees.

### Metagenomic analyses

675 metagenomes were obtained from TARA and BioGeoTRACES (76,77). These metagenomes were searched for SAR11 purine usage genes and SAR11 ribosomal proteins using custom Hidden Markov Models (HMM’s) that were created as follows. First, a set of ∼20 representative sequences for the genes of each individual step in the purine catabolism pathway (Figs. 3,4), as well as for a set of 14 single-copy ribosomal proteins (78) were obtained at Uniprot (79) and assembled into HMM’s. For phosphoribosyltransferase, which has many homologs operating on different nucleobases, the initial set was very narrowly chosen to consist of primarily SAR11 variants occurring with purine catabolism operons. Resulting HMM’s were searched against the Uniprot database using the PHMMER algorithm within the HMMER web portal (80), and results were manually inspected to determine e-value cut offs (Data S3) that captured maximal diversity while eliminating homologous genes associated with other functions. The original 20-gene HMM’s were then searched against the MARMICRODB database (33,74) and all sequences falling below the e-value cutoffs determined from HMMER results were included our final HMM’s to ensure they have broad coverage of diversity within the extant microbial oceans. Next, our custom HMM’s were searched against our full set of metagenomes using the GraftM algorithm (78). The results of this search were again manually inspected to determine GraftM cutoff scores (Data S3) that mostly eliminate homologous genes associated with other functions while mostly capturing sequences of interest. Taxonomy of metagenome reads that mapped to our HMM’s was assigned using the pplacer function, which places reads in a phylogenetic tree implicitly generated from the sequence alignment underlying an HMM (81). To estimate gene/genome frequencies of individual purine usage genes in SAR11, the counts for each purine usage gene were then normalized to the HMM amino acid length and to the median count of the 14 single-copy ribosomal protein genes. Similarity in the resulting biogeographical distributions of genes grouped together into overarching purine usage functions (Fig. S4-S9), each of which were independently analyzed, confirms robustness of the approach.

Chemical and hydrographic data associated with the BioGEOTRACES metagenomes (77) were obtained from the GEOTRACES Intermediate Data Product IDP2017 version 2 (accessed January 2019), specifically from sections GA02 (82,83) GA03, GA10 (82), and GP13. Environmental data from the Tara Oceans project was obtained from https://doi.pangaea.de/10.1594/PANGAEA.875579. Metagenome sequence data were quality controlled, environmental data matched to metagenomic samples, and missing environmental data were imputed as described in detail in the supplementary material of Hogle et al. 2022 (85). Spearman-rank correlations between nutrient concentrations and gene frequencies were determined in R.

Purine usage genes were grouped into overarching functional categories using a combination of their biogeography (Fig. S5-S9), correlations with nutrient concentrations (Fig. S4, Table S2), and correlations in their gene/genome frequencies across metagenomes (Table S6). We assumed that frequencies of genes would show greater correlations to those of genes belonging to the same functional categories than for genes belonging to different categories, and that we could therefore use correlations as one criterion for grouping genes. To this end, Pearson correlation coefficients for the metagenomic gene/genome frequencies of all genes relative to each other were determined in Excel (Table S6). The frequency of one SAR11 purine deaminase homolog (here “purine deaminase 2”), whose biochemical function involves nitrogen harvesting, was uncorrelated with nutrient concentrations (Fig. S4, Table S2), whereas the frequencies of other genes associated with nitrogen harvesting are negatively correlated with nutrient concentrations (Fig. S4, Table S2). However, the frequency of the purine deaminase 2 gene was highly correlated with the frequencies of other genes involved in nitrogen harvesting (Table S6), and like other genes in this category remained below ∼40-45% in our full dataset, leading us to conclude they group together.

### Metatranscriptome analysis

Time-resolved in situ community gene expression data from the North Pacific were obtained from Ref (51). Transcript counts for individual genes were normalized to total SAR11 or *Prochlorococcus* transcripts at each timepoint, and results were further z-score normalized to allow comparison across genes (Data S2).

### SAR11 adenine amendment experiments

Cells of SAR11 strain HTCC7211 were cultured under constant light at 22 °C in ProMS medium (33). ProMS is a seawater-based modified version of the artificial AMS1 medium (15), in which the concentrations of the main sources of carbon (pyruvate), nitrogen (glycine) and sulfur (methionine) have been lowered 50-fold to promote detection of the impact of added organic substrates or phytoplankton exudates (33). In addition, HTCC7211 cells were cultured in a glycine-depleted version of ProMS in which the concentration of all nutrients other than glycine was increased 50-fold to bring them back to the levels of the AMS1 medium. These cultures were then amended with adenine at concentrations ranging seven orders of magnitude, from 10 pM to 100 µM, and their growth was tracked alongside that of adenine-free control cultures for an entire growth curve.

Culture density and purity and cell size and DNA content were measured using flow cytometry. Because sub-populations in different phases of the cell cycle are difficult to distinguish in SAR11 (15), we instead quantified the averages and standard deviations over the total bulk fluorescence distributions (Fig. S12). To examine whether adenine exposure poises SAR11 cells for replication, cultures grown with and without adenine were harvested in mid-exponential growth and washed three times using adenine-free culture media and ultracentrifugation at ∼25,000 x g (15,000 rpm) for 45 minutes at 22 °C. After the final washing step cell pellets were resuspended in adenine-free media and subsequent growth was monitored via flow cytometry. Fisher’s exact tests of the significance in the differences of the number of cultures experiencing lags were determined in R.

## Acknowledgments

We thank Eli Salcedo for help in generating relative DNA fluorescence/cell histograms from raw flow cytometry data for SAR11 adenine amendment experiments, Gretchen Swarr for help in the processing of samples for metabolomics analysis, and Steve Biller for discussions regarding genome replication in bacteria. This research was supported by grants from the Simons Foundation (Award ID 509034SCFY20 to R.B. and S.W.C., Award ID 509034FY20 to E.B.K., and SCOPE Award ID 329108 to M.J. Follows) and from the National Science Foundation (OCE-2019589 to R.B.). This is C-CoMP publication #33.

## Author contributions

Conceptualization: RB

Methodology: RB, BS, TJO, KL, SLH, JWB, RCL, KD, AA, MCKS, EBK

Investigation: RB, BS, TJO, KL, JWB, RCL, KD, AA, MCKS

Formal Analysis, RB, BS, TJO, KL, SLH, RCL, MCKS

Visualization: RB, BS

Funding acquisition: RB, SWC, EBK

Writing—original draft: RB

Writing—review & editing: RB, BS, TJO, KL, SLH, JWB, RCL, KD, AA, MCKS, EBK, SWC

## Data and materials availability

All raw metabolomic, genomic or gene expression data used in this work come from previous studies and are publicly available as described in the Methods. All other data, including those resulting from the analysis of previously existing data, are available in the main text or the supplementary materials.

## References

1. L.R. Pomeroy, The ocean’s food web, a changing paradigm. Bioscience, 24(9), pp.499–504 (1974)

2. F. Azam, T. Fenchel, J.G. Field, J.S. Gray, L.A. Meyer-Reil, F. Thingstad, The ecological role of water-column microbes in the sea. Marine ecology progress series. Oldendorf, 10(3), pp.257–263 (1983)

3. M.A. Moran, E.B. Kujawinski, W.F. Schroer, S.A. Amin, N.R. Bates, E.M. Bertrand, R. Braakman, C.T. Brown, M.W. Covert, S.C. Doney, S.T. Dyhrman, A.S. Edison, A.M. Eren, N.M. Levine, L. Li, A.C. Ross, M.A. Saito, A.E. Santoro, D. Segrè, A. Shade, M.B. Sullivan, A. Vardi, Microbial metabolites in the marine carbon cycle. Nature microbiology, 7(4), pp.508–523 (2022)

4. C.B. Field, M.J. Behrenfeld, J.T. Randerson, P. Falkowski, Primary production of the biosphere: integrating terrestrial and oceanic components. science, 281(5374), pp. 237–240 (1998)

5. P.G. Falkowski, J.A. Raven, Aquatic photosynthesis. Princeton University Press (2013)

6. R. Braakman, E. Smith, The compositional and evolutionary logic of metabolism. Physical biology, 10(1), p. 011001 (2012)

7. J.A. Fuhrman, Marine viruses and their biogeochemical and ecological effects. Nature, 399(6736), pp.541–548 (1999)

8. S.W. Wilhelm, C.A. Suttle, Viruses and nutrient cycles in the sea: viruses play critical roles in the structure and function of aquatic food webs. Bioscience, 49(10), pp.781–788 (1999)

9. B.A. Ward, S. Dutkiewicz, O. Jahn, M.J. Follows, A size-structured food-web model for the global ocean. Limnology and Oceanography, 57(6), pp.1877–1891 (2012)

10. B.P. Durham, S. Sharma, H. Luo, C.B. Smith, S.A. Amin, S.J. Bender, S.P. Dearth, B.A. Van Mooy, S.R. Campagna, E.B. Kujawinski, E.V. Armbrust, Cryptic carbon and sulfur cycling between surface ocean plankton. Proceedings of the National Academy of Sciences, 112(2), pp.453–457 (2015)

11. E. Segev, T.P. Wyche, K.H, Kim, J. Petersen, C. Ellebrandt, H. Vlamakis, N. Barteneva, J.N. Paulson, L. Chai, J. Clardy, R. Kolter, Dynamic metabolic exchange governs a marine algal-bacterial interaction. elife, 5, p.e17473 (2016)

12. R.K. Fritts, A.L. McCully, J.B. McKinlay, Extracellular metabolism sets the table for microbial cross-feeding. Microbiology and Molecular Biology Reviews, 85(1), pp.e00135–20 (2021)

13. S. Giri, L. Oña, S. Waschina, S. Shitut, G. Yousif, C. Kaleta, C. Kost, Metabolic dissimilarity determines the establishment of cross-feeding interactions in bacteria. Current Biology, 31(24), pp.5547–5557 (2021)

14. S.J. Giovannoni, H.J. Tripp, S. Givan, M. Podar, K.L. Vergin, D. Baptista, L. Bibbs, J. Eads, T.H. Richardson, M. Noordewier, M.S. Rappé, Genome streamlining in a cosmopolitan oceanic bacterium. Science, 309(5738), pp.1242–1245 (2005)

15. P. Carini, L. Steindler, S. Beszteri, S.J. Giovannoni, Nutrient requirements for growth of the extreme oligotroph ‘Candidatus Pelagibacter ubique’HTCC1062 on a defined medium. The ISME journal, 7(3), pp.592–602 (2013)

16. B.K. Swan, B. Tupper, A. Sczyrba, F.M. Lauro, M. Martinez-Garcia, J.M. González, H. Luo, J.J. Wright, Z.C. Landry, N.W. Hanson, B.P. Thompson, Prevalent genome streamlining and latitudinal divergence of planktonic bacteria in the surface ocean. Proceedings of the National Academy of Sciences, 110(28), pp.11463–11468 (2013)

17. S.W. Chisholm, R.J. Olson, E.R. Zettler, R. Goericke, J.B. Waterbury, J.B. N.A. Welschmeyer, A novel free-living prochlorophyte abundant in the oceanic euphotic zone. Nature, 334(6180), pp.340–343 (1988)

18. P. Flombaum, J.J. Gallegos, R.A. Gordillo, J. Rincón, L.L. Zabala, N. Jiao, D.M. Karl, W.K. Li, M.W. Lomas, D. Veneziano, C.S. Vera, Present and future global distributions of the marine Cyanobacteria Prochlorococcus and Synechococcus. Proceedings of the National Academy of Sciences, 110(24), pp.9824–9829 (2013)

19. R.T. Letscher, J.K. Moore, A.C. Martiny, M.W. and Lomas, Biodiversity and Stoichiometric Plasticity Increase Pico-Phytoplankton Contributions to Marine Net Primary Productivity and the Biological Pump. Global Biogeochemical Cycles, 37(8), p.e2023GB007756 (2023)

20. R. Braakman, M.J. Follows, S.W. Chisholm, Metabolic evolution and the self-organization of ecosystems. Proceedings of the National Academy of Sciences, 114(15), pp.E3091–E3100 (2017)

21. E.B. Kujawinski, R. Braakman, K. Longnecker, J.W. Becker, S.W. Chisholm, K. Dooley, M.C. Kido Soule, G.J. Swarr, K. Halloran, Metabolite diversity among Prochlorococcus strains belonging to divergent ecotypes. mSystems, e01261–22 (2023)

22. A. Dufresne, L. Garczarek, F. Partensky, Accelerated evolution associated with genome reduction in a free-living prokaryote. Genome biology, 6, pp.1–10 (2005)

23. J.J. Grzymski, A.M. Dussaq, The significance of nitrogen cost minimization in proteomes of marine microorganisms. The ISME journal, 6(1), pp.71–80 (2012)

24. M. Proudfoot, E. Kuznetsova, G. Brown, N.N. Rao, M. Kitagawa, H. Mori, A. Savchenko, A.F. Yakunin, General enzymatic screens identify three new nucleotidases in Escherichia coli: biochemical characterization of SurE, YfbR, and YjjG. Journal of Biological Chemistry, 279(52), pp.54687–54694 (2004)

25. M. Vodnala, F. Ranjbarian, A. Pavlova, H.P. de Koning, A. Hofer, Trypanosoma brucei methylthioadenosine phosphorylase protects the parasite from the antitrypanosomal effect of deoxyadenosine: implications for the pharmacology of adenosine antimetabolites. Journal of Biological Chemistry, 291(22), pp.11717–11726 (2016)

26. G. Cacciapuoti, C. Bertoldo, A. Brio, V. Zappia, M. Porcelli, Purification and characterization of 5′-methylthioadenosine phosphorylase from the hyperthermophilic archaeon Pyrococcus furiosus: substrate specificity and primary structure analysis. Extremophiles, 7, pp.159–168 (2003)

27. S.E. Noell, S.J. Giovannoni, SAR11 bacteria have a high affinity and multifunctional glycine betaine transporter. Environmental Microbiology, 21(7), pp.2559–2575 (2019)

28. S.E. Noell, G.E. Barrell, C. Suffridge, J. Morré, K.P. Gable, J.R. Graff, B.J. VerWey, F.L. Hellweger, S.J. Giovannoni, Sar11 cells rely on enzyme multifunctionality to metabolize a range of polyamine compounds. MBio, 12(4), pp.e01091–21 (2021)

29. E.R. Zinser, D. Lindell, Z.I. Johnson, M.E. Futschik, C. Steglich, M.L. Coleman, M.A. Wright, T. Rector, R. Steen, N. McNulty, L.R. Thompson, Choreography of the transcriptome, photophysiology, and cell cycle of a minimal photoautotroph, Prochlorococcus. PloS one, 4(4), p.e5135 (2009)

30. Y.C. Chuang, N.W. Haas, R. Pepin, M.G. Behringer, Y. Oda, B. LaSarre, C.S. Harwood, J.B. McKinlay, Bacterial adenine cross-feeding stems from a purine salvage bottleneck. The ISME Journal, 18(1), p.wrae034 (2024)

31. R.M. Morris, M.S. Rappé, S.A. Connon, K.L. Vergin, W.A. Siebold, C.A. Carlson, S.J. Giovannoni, SAR11 clade dominates ocean surface bacterioplankton communities. Nature, 420(6917), pp.806–810 (2002)

32. A.H. Treusch, K.L. Vergin, L.A. Finlay, M.G. Donatz, R.M. Burton, C.A. Carlson, S.J. Giovannoni, Seasonality and vertical structure of microbial communities in an ocean gyre. The ISME journal, 3(10), pp.1148–1163 (2009)

33. J.W. Becker, S.L. Hogle, K. Rosendo, S.W. Chisholm, Co-culture and biogeography of Prochlorococcus and SAR11. The ISME journal, 13(6), pp.1506–1519 (2019)

34. J.M. Haro-Moreno, F. Rodriguez-Valera, R. Rosselli, F. Martinez-Hernandez, J.J. Roda-Garcia, M.L. Gomez, O. Fornas, M. Martinez-Garcia, M. López-Pérez, Ecogenomics of the SAR11 clade. Environmental Microbiology, 22(5), pp.1748–1763 (2020)

35. N.J. Antia, B.R. Berland, D.J. Bonin, S.Y. Maestrini, Allantoin as nitrogen source for growth of marine benthic microalgae. Phycologia, 19(2), pp.103–109 (1980)

36. J.F. Weber, F.A. Fuhrman, G.J. Fuhrman, H.S. Mosher, Isolation of allantoin and adenosine from the marine sponge Tethya aurantia. Comparative Biochemistry and Physiology Part B: Comparative Biochemistry, 70(4), pp.799–801 (1981)

37. G.V.D. Vogels, C. Van der Drift, Degradation of purines and pyrimidines by microorganisms. Bacteriological reviews, 40(2), pp.403–468 (1976)

38. A.K. Werner, C.P. Witte, The biochemistry of nitrogen mobilization: purine ring catabolism. Trends in Plant Science, 16(7), pp.381–387 (2011)

39. S. Watanabe, M. Matsumoto, Y. Hakomori, H. Takagi, H. Shimada, A. Sakamoto, The purine metabolite allantoin enhances abiotic stress tolerance through synergistic activation of abscisic acid metabolism. Plant, cell & environment, 37(4), pp.1022–1036 (2014)

40. S.J. Giovannoni, SAR11 bacteria: the most abundant plankton in the oceans. Annual review of marine science, 9, pp.231–255 (2017)

41. L.R. Moore, A.F. Post, G. Rocap, S.W. Chisholm, Utilization of different nitrogen sources by the marine cyanobacteria Prochlorococcus and Synechococcus. Limnology and oceanography, 47(4), pp.989–996 (2002)

42. H. Berthelot, S. Duhamel, S. L’helguen, J.F. Maguer, S. Wang, I. Cetinić, N. Cassar, NanoSIMS single cell analyses reveal the contrasting nitrogen sources for small phytoplankton. The ISME Journal, 13(3), pp.651–662 (2019)

43. C.L. Fiore, K. Longnecker, M.C. Kido Soule, E.B. Kujawinski, Release of ecologically relevant metabolites by the cyanobacterium *Synechococcus elongatus* CCMP 1631. Environmental microbiology, 17(10), pp.3949–3963 (2015)

44. M.B. Karner, E.F. DeLong, D.M. Karl, Archaeal dominance in the mesopelagic zone of the Pacific Ocean. Nature, 409(6819), pp.507–510 (2001)

45. B. Bayer, R.L. Hansman, M.J. Bittner, B.E. Noriega-Ortega, J. Niggemann, T. Dittmar, G.J. Herndl, Ammonia-oxidizing archaea release a suite of organic compounds potentially fueling prokaryotic heterotrophy in the ocean. Environmental Microbiology, 21(11), pp.4062–4075 (2019)

46. K. Kitzinger, C.C. Padilla, H.K. Marchant, P.F. Hach, C.W. Herbold, A.T. Kidane, M. Könneke, S. Littmann, M. Mooshammer, J. Niggemann, S. Petrov, Cyanate and urea are substrates for nitrification by Thaumarchaeota in the marine environment. Nature microbiology, 4(2), pp.234–243 (2019)

47. J.C. Thrash, B. Temperton, B.K. Swan, Z.C. Landry, T. Woyke, E.F. DeLong, R. Stepanauskas, S.J. Giovannoni, Single-cell enabled comparative genomics of a deep ocean SAR11 bathytype. The ISME journal, 8(7), pp.1440–1451 (2014)

48. D.C. Rees, E. Johnson, O. Lewinson, ABC transporters: the power to change. Nature reviews Molecular cell biology, 10(3), pp.218–227 (2009)

49. S. Weyand, T. Shimamura, S. Yajima, S.I. Suzuki, O. Mirza, K. Krusong, E.P. Carpenter, N.G. Rutherford, J.M. Hadden, J. O’Reilly, P. Ma, Structure and molecular mechanism of a nucleobase– cation–symport-1 family transporter. Science, 322(5902), pp.709–713 (2008)

50. D. Muratore, A.K. Boysen, M.J. Harke, K.W. Becker, J.R. Casey, S.N. Coesel, D.R. Mende, S.T. Wilson, F.O. Aylward, J.M. Eppley, A. Vislova, Complex marine microbial communities partition metabolism of scarce resources over the diel cycle. Nature Ecology & Evolution, 6(2), pp.218–229 (2022)

51. E.A. Ottesen, C.R. Young, S.M. Gifford, J.M. Eppley, R. Marin III, S.C. Schuster, C.A. Scholin, E.F. DeLong, Multispecies diel transcriptional oscillations in open ocean heterotrophic bacterial assemblages. Science, 345(6193), pp.207–212 (2014)

52. R.H. MacArthur, Population ecology of some warblers of northeastern coniferous forests. Ecology, 39(4), pp.599–619 (1958)

53. M.L. Coleman, S.W. Chisholm, Ecosystem-specific selection pressures revealed through comparative population genomics. Proceedings of the National Academy of Sciences, 107(43), pp.18634–18639 (2010)

54. L.J. Ustick, A.A. Larkin, C.A. Garcia, N.S. Garcia, M.L. Brock, J.A. Lee, N.A. Wiseman, J.K. Moore, A.C. Martiny, Metagenomic analysis reveals global-scale patterns of ocean nutrient limitation. Science, 372(6539), pp.287–291 (2021)

55. M. Acker, S.L. Hogle, P.M. Berube, T. Hackl, A. Coe, R. Stepanauskas, S.W. Chisholm, D.J. Repeta, Phosphonate production by marine microbes: exploring new sources and potential function. Proceedings of the National Academy of Sciences, 119(11), p.e2113386119 (2022)

56. W.F. Doolittle, A. Booth, It’s the song, not the singer: an exploration of holobiosis and evolutionary theory. Biology & Philosophy, 32, pp.5–24 (2017)

57. C.A. Carlson, R. Morris, R. Parsons, A.H. Treusch, S.J. Giovannoni, K. Vergin, Seasonal dynamics of SAR11 populations in the euphotic and mesopelagic zones of the northwestern Sargasso Sea. The ISME journal, 3(3), pp.283–295 (2009)

58. M.V. Brown, F.M. Lauro, M.Z. DeMaere, L. Muir, D. Wilkins, T. Thomas, M.J. Riddle, J.A. Fuhrman, C. Andrews-Pfannkoch, J.M. Hoffman, J.B. McQuaid, Global biogeography of SAR11 marine bacteria. Molecular systems biology, 8(1), p.595 (2012)

59. A. Eiler, R. Mondav, L. Sinclair, L. Fernandez-Vidal, D.G. Scofield, P. Schwientek, M. Martinez-Garcia, D. Torrents, K.D. McMahon, S.G. Andersson, R. Stepanauskas, Tuning fresh: radiation through rewiring of central metabolism in streamlined bacteria. The ISME journal, 10(8), pp.1902–1914 (2016)

60. E.A. Ottesen, C.R. Young, J.M. Eppley, J.P. Ryan, F.P. Chavez, C.A. Scholin, E.F. DeLong, Pattern and synchrony of gene expression among sympatric marine microbial populations. Proceedings of the National Academy of Sciences, 110(6), pp.E488–E497 (2013)

61. B.C. Goodwin, Oscillatory behavior in enzymatic control processes. Advances in enzyme regulation, 3, pp.425–437 (1965)

62. D. Gonze, J. Halloy, A. Goldbeter, Robustness of circadian rhythms with respect to molecular noise. Proceedings of the National Academy of Sciences, 99(2), pp.673–678 (2002)

63. B. Novák, J.J. Tyson, Design principles of biochemical oscillators. Nature reviews Molecular cell biology, 9(12), pp.981–991 (2008)

64. N. Xeros, Deoxyriboside control and synchronization of mitosis. Nature, 194(4829), pp.682–683 (1962)

65. J.M. Mitchison, J. Creanor, Induction synchrony in the fission yeast Schizosaccharomyces pombe. Experimental cell research, 67(2), pp.368–374 (1971)

66. G. Bjursell, P. Reichard, Effects of thymidine on deoxyribonucleoside triphosphate pools and deoxyribonucleic acid synthesis in Chinese hamster ovary cells. Journal of Biological Chemistry, 248(11), pp.3904–3909 (1973)

67. A. Jordan, P. Reichard, Ribonucleotide reductases. Annual review of biochemistry, 67(1), pp.71–98 (1998)

68. A.A. Larkin, G.I. Hagstrom, M.L. Brock, N.S. Garcia, A.C. Martiny, Basin-scale biogeography of Prochlorococcus and SAR11 ecotype replication. The ISME journal, 17(2), pp.185–194 (2023)

69. W.M. Johnson, M.C. Kido Soule, E.B. Kujawinski, Extraction efficiency and quantification of dissolved metabolites in targeted marine metabolomics. Limnology and Oceanography: Methods, 15(4), pp.417–428 (2017)

70. C.S. Ting, C. Hsieh, S. Sundararaman, C. Mannella, M. Marko, Cryo-electron tomography reveals the comparative three-dimensional architecture of Prochlorococcus, a globally important marine cyanobacterium. Journal of bacteriology, 189(12), pp.4485–4493 (2007)

71. N. Cermak, J.W. Becker, S.M. Knudsen, S.W. Chisholm, S.R. Manalis, M.F. Polz, Direct single-cell biomass estimates for marine bacteria via Archimedes’ principle. The ISME journal, 11(3), pp.825–828 (2017)

72. M. Kanehisa, M. Furumichi, M. Tanabe, Y. Sato, K. Morishima, KEGG: new perspectives on genomes, pathways, diseases and drugs. Nucleic acids research, 45(D1), pp.D353–D361 (2017)

73. M.Z. Cader, R.P. de Almeida Rodrigues, J.A. West, G.W. Sewell, N. Muhammad, S. Reikine, G. Sirago, L.W. Unger, A.B. Iglesias-Romero, K. Ramshorn, L.M. Haag, FAMIN is a multifunctional purine enzyme enabling the purine nucleotide cycle. Cell, 180(2), pp.278–295 (2020)

74. S.L. Hogle, MARMICRODB database for taxonomic classification of (marine) metagenomes (1.0.0) [Data set]. Zenodo. 10.5281/zenodo.3520509 (2019)

75. C. Camacho, G. Coulouris, V. Avagyan, N. Ma, J. Papadopoulos, K. Bealer, T.L. Madden, BLAST+: architecture and applications. BMC bioinformatics, 10, pp.1–9 (2009)

76. S. Sunagawa, et al., Structure and function of the global ocean microbiome. Science, 348(6237), p.1261359 (2015)

77. S.J. Biller, P.M. Berube, K. Dooley, M. Williams, B.M. Satinsky, T. Hackl, S.L. Hogle, A. Coe, K. Bergauer, H.A. Bouman, T.J. Browning, Marine microbial metagenomes sampled across space and time. Scientific data, 5(1), pp.1–7 (2018)

78. J.A. Boyd, B.J. Woodcroft, G.W. Tyson, GraftM: a tool for scalable, phylogenetically informed classification of genes within metagenomes. Nucleic Acids Research, 46(10), pp.e59–e59 (2018)

79. The Uniprot Consortium, UniProt: the universal protein knowledgebase in 2023. Nucleic Acids Research 51, no. D1: D523–D531 (2023)

80. S.C. Potter, A. Luciani, S.R. Eddy, Y. Park, R. Lopez, R.D. Finn, HMMER web server: 2018 update. Nucleic acids research, 46(W1), pp.W200–W204 (2018)

81. F.A. Matsen, R.B. Kodner, E.V. Armbrust, pplacer: linear time maximum-likelihood and Bayesian phylogenetic placement of sequences onto a fixed reference tree. BMC bioinformatics, 11, pp.1–16 (2010)

82. L.A. Salt, S.M. van Heuven, M.E. Claus, E.M. Jones, H.J.W. De Baar, Rapid acidification of mode and intermediate waters in the southwestern Atlantic Ocean. Biogeosciences, 12(5), pp.1387–1401 (2015)

83. M.J. Rijkenberg, R. Middag, P. Laan, L.J. Gerringa, H.M. Aken, V. Schoemann, J.T. de Jong, H.J. De Baar, The distribution of dissolved iron in the West Atlantic Ocean. PloS one, 9(6), p.e101323 (2014)

84. N.J. Wyatt, A. Milne, E.M.S. Woodward, A.P. Rees, T.J. Browning, H.A. Bouman, P.J. Worsfold, M.C. Lohan, Biogeochemical cycling of dissolved zinc along the GEOTRACES South Atlantic transect GA10 at 40 S. Global Biogeochemical Cycles, 28(1), pp.44–56 (2014)

85. S.L. Hogle, T. Hackl, R.M. Bundy, J. Park, B. Satinsky, T. Hiltunen, S. Biller, P.M. Berube, S.W. Chisholm, Siderophores as an iron source for picocyanobacteria in deep chlorophyll maximum layers of the oligotrophic ocean. The ISME Journal, 16(6), pp.1636–1646 (2022)

