## Supplementary Information for "Global niche partitioning of purine and pyrimidine cross-feeding among ocean microbes"

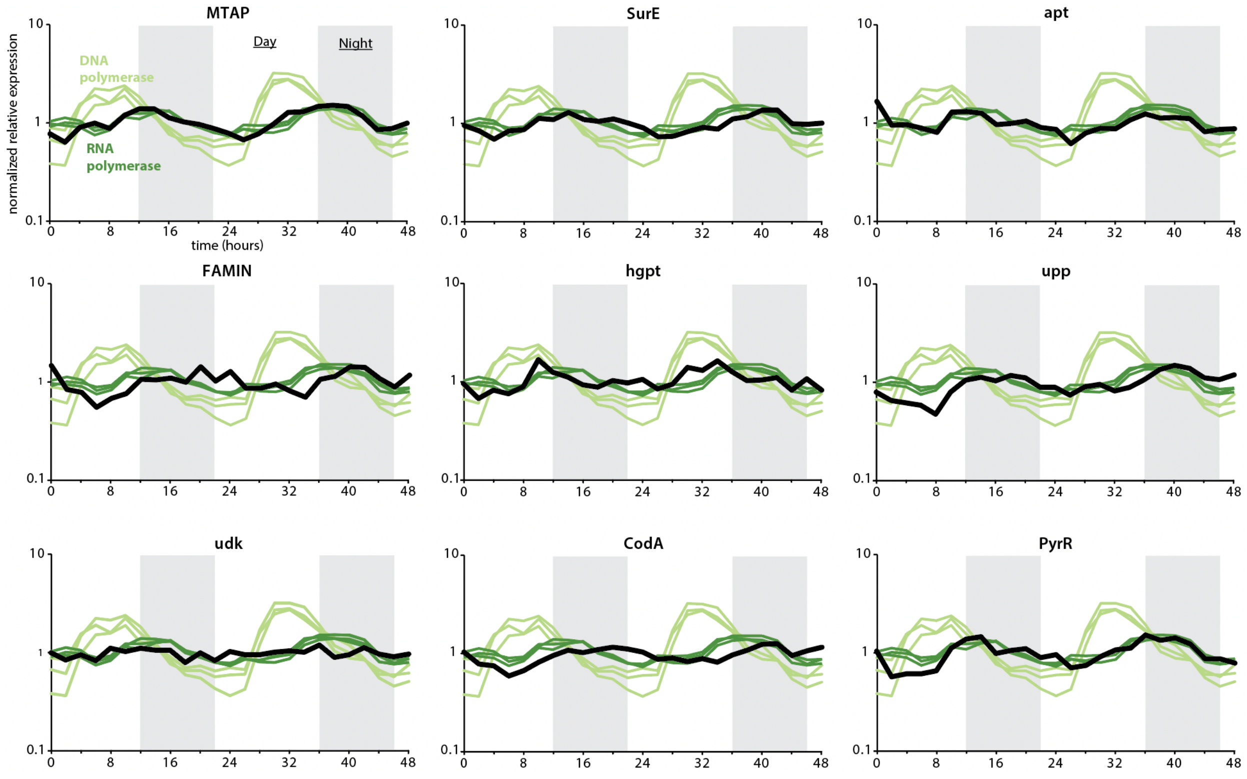

**Fig. S1. Transcription dynamics of possible genes involved in (deoxy)ribonucleotide recycling in *Prochlorococcus*.** Data was obtained from Ref. (29) for a *Prochlorococcus* strain (MED4) whose growth was synchronized to diurnal L:D conditions. Focal genes are in black in each panel, whereas DNA and RNA polymerase genes are in light and dark green, respectively.

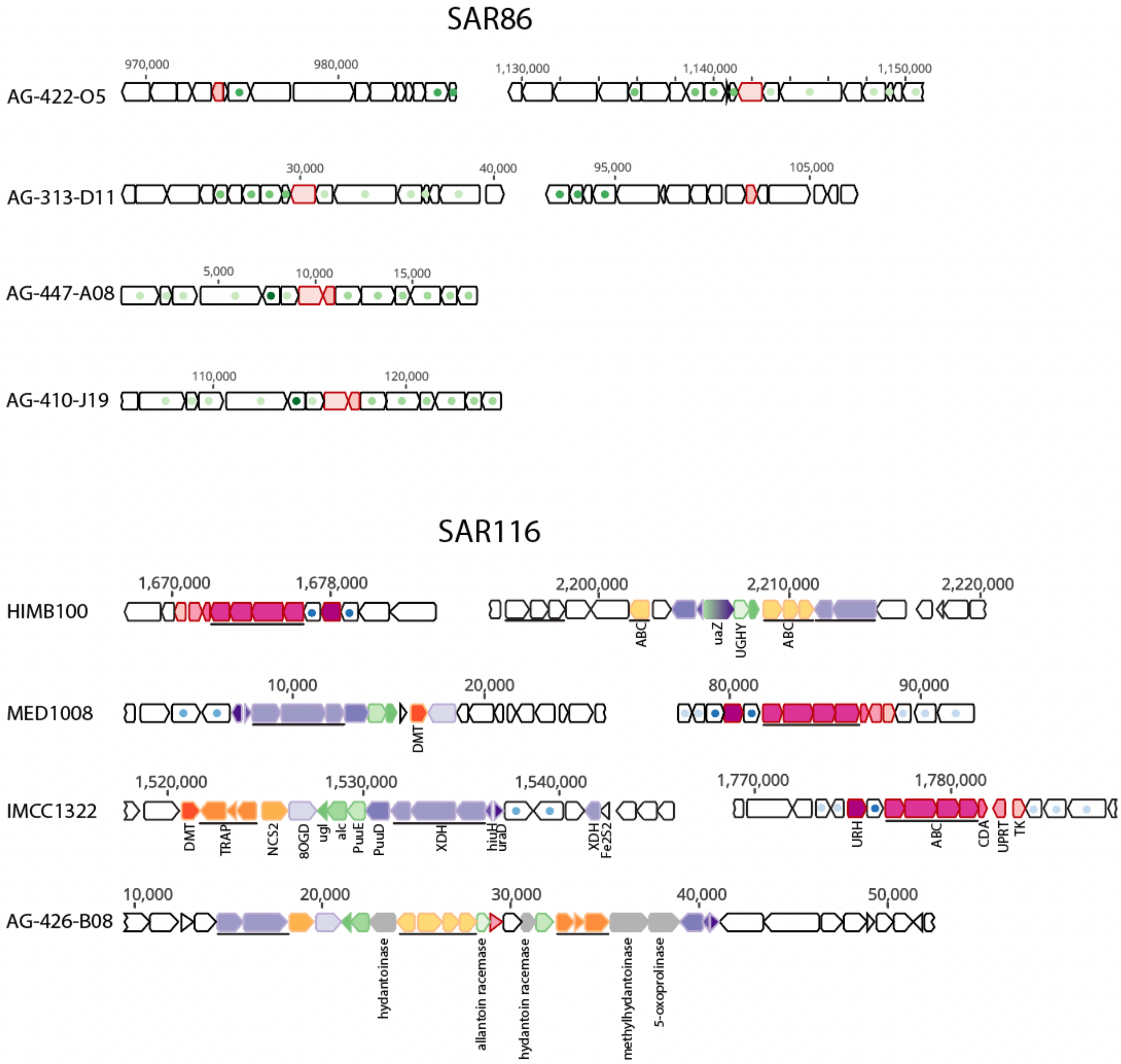

**Fig. S2. Pyrimidine and purine usage genes in representative genomes of SAR86 and SAR116.** Pyrimidine usage genes are in shades of pink, purine assimilation genes are in shades of yellow-orange, purine catabolism genes involved in energy harvesting are in shades of purple and allantoin catabolism genes are in shades of green. Genes with colored dots are conserved flanking genes.

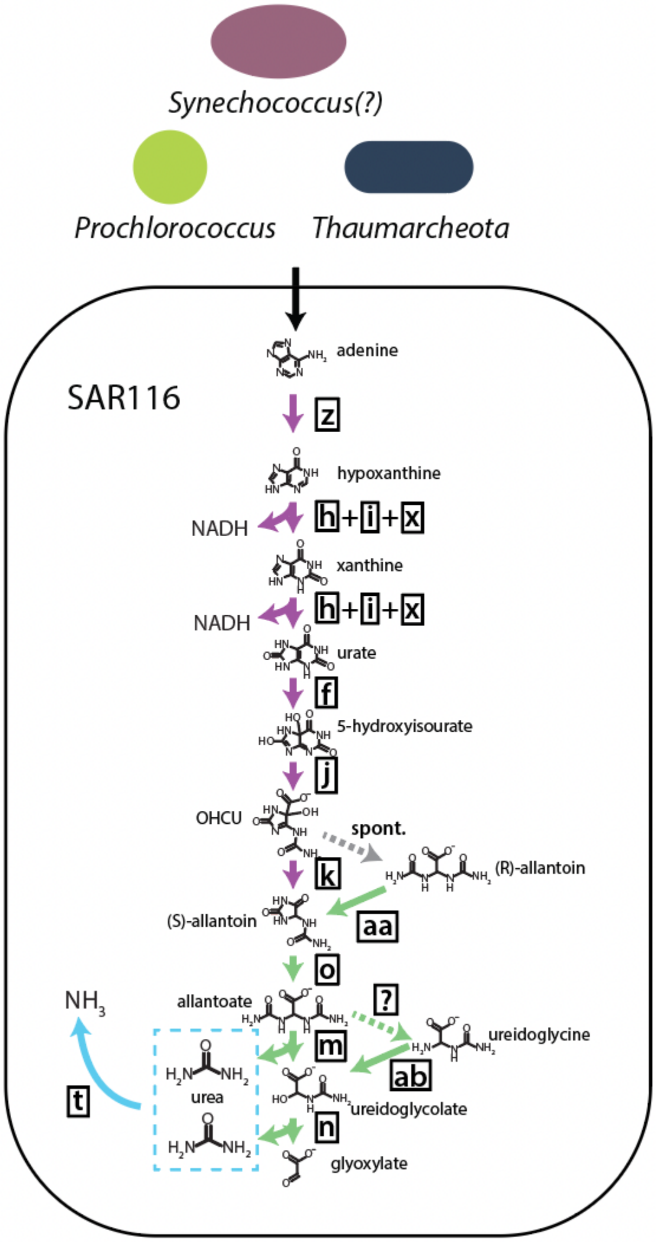

**Fig. S3. Variants of purine catabolism in SAR116.** *Prochlorococcus, Thaumarcheota* and potentially *Synechococcus* (see main text) supply purines that are catabolized by SAR116. Reactions that perform different purine usage functions represented in different colors as defined in Fig. 3. Some genomes contain genes for allantoin racemase (aa) and/or ureidoglycine aminohydrolase (ab) instead of allantoicase (m), suggesting variations in how SAR116 process purines. Molecule names are listed below or to the right or left of molecules, while boxed letters refer to enzymes as listed in Fig. 3. Black arrows represent inferred cross-feeding pathways. Dashed arrows mark a spontaneous reaction (gray) and a reaction for which enzymes were not identified (green), with the latter also marked with a boxed question remark representing the unidentified enzyme.

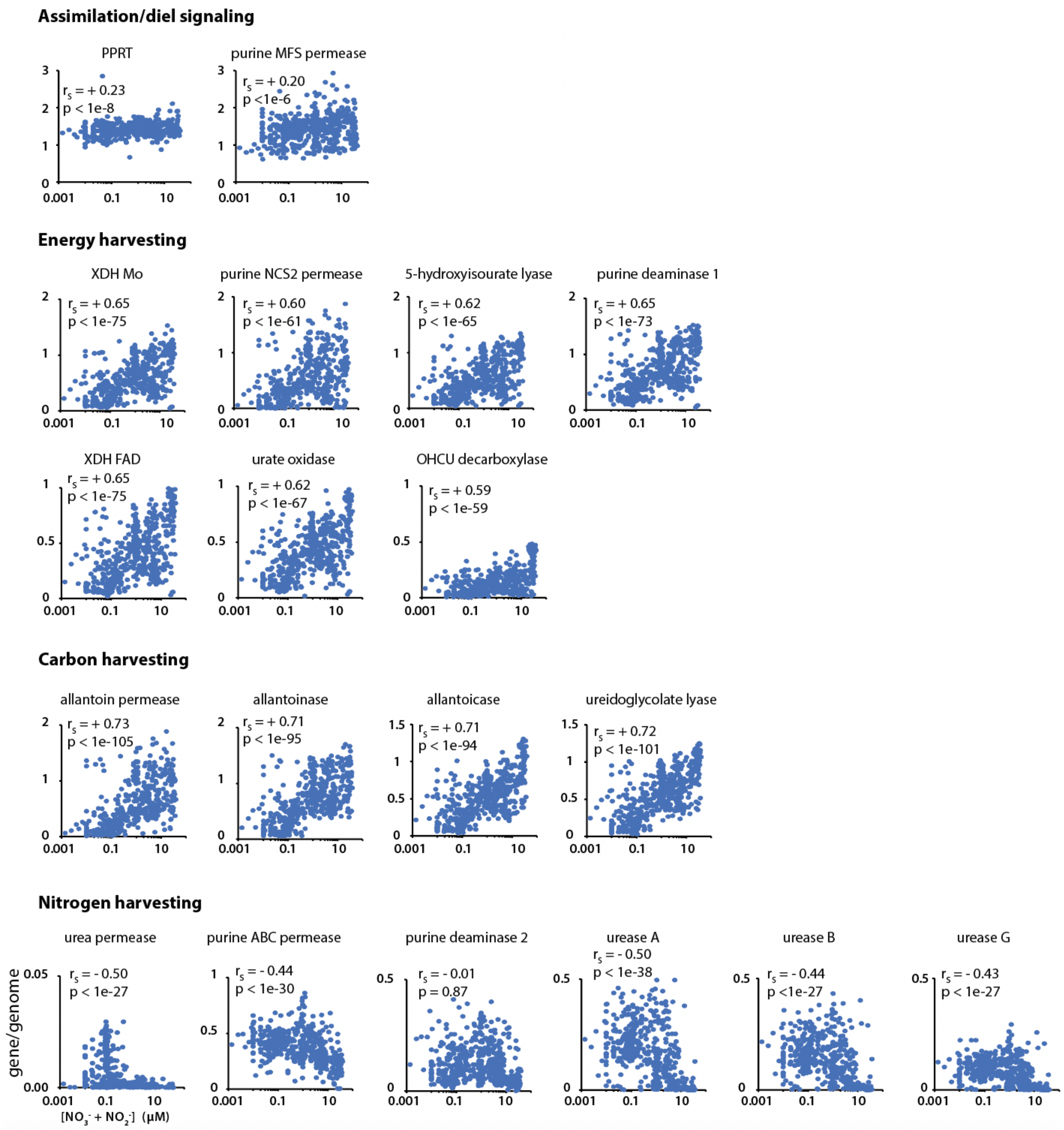

**Fig. S4. Frequencies of purine usage genes per SAR11 genome as a function of nitrate and nitrate concentrations in TARA and BioGEOTRACES metagenomes.** Gene frequencies are normalized to frequencies of single-copy ribosomal proteins to give an estimate of gene per SAR11 genome. Spearman rank coefficients ($r_S$) and associated p-value of the correlation between gene frequencies and nitrate and nitrate concentrations (see Table S2) are shown for each gene.

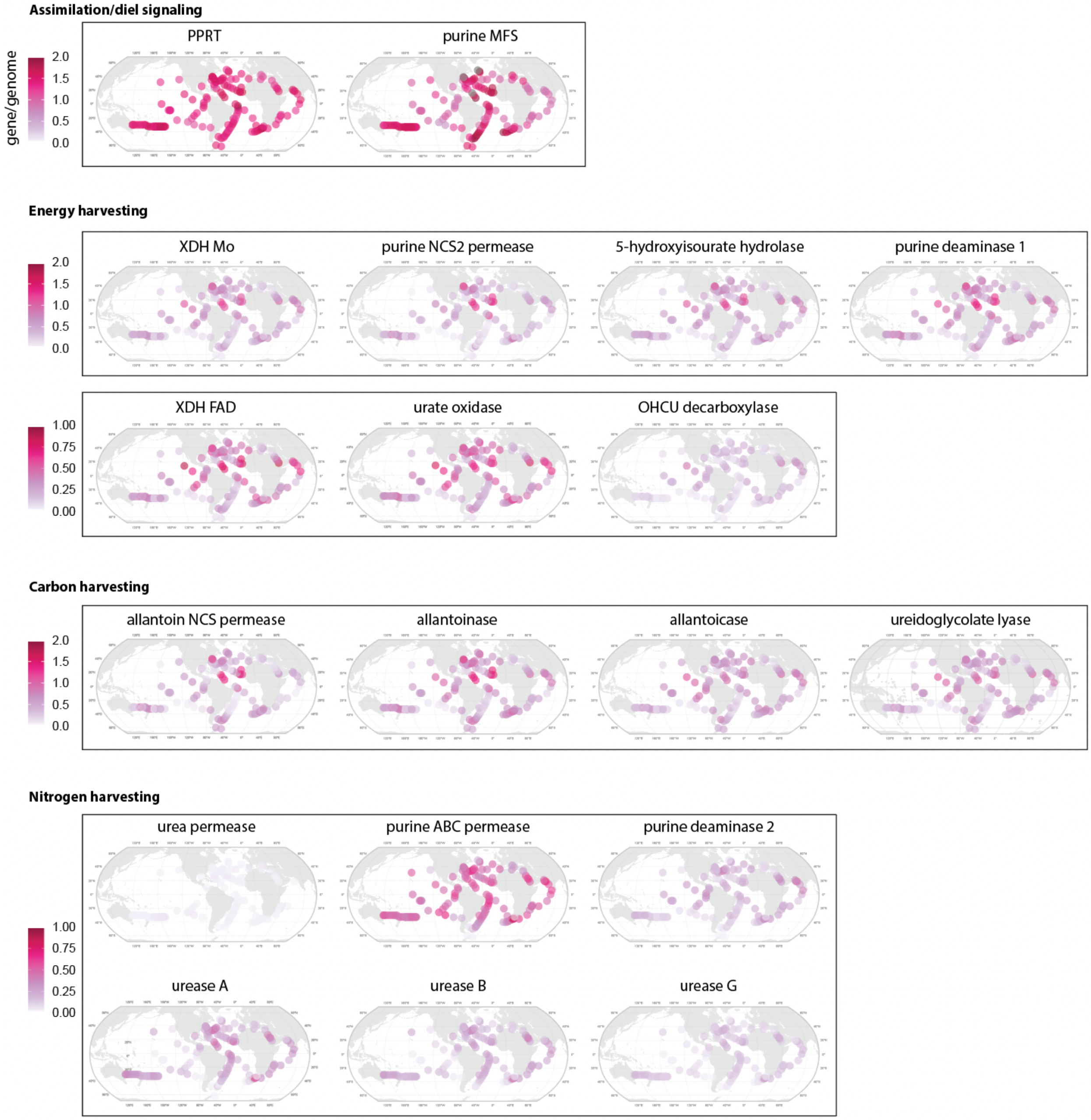

**Fig. S5. Biogeographical distribution of frequency of purine usage genes per SAR11 genome.** Global distributions are within the upper ~100 meters of water column.

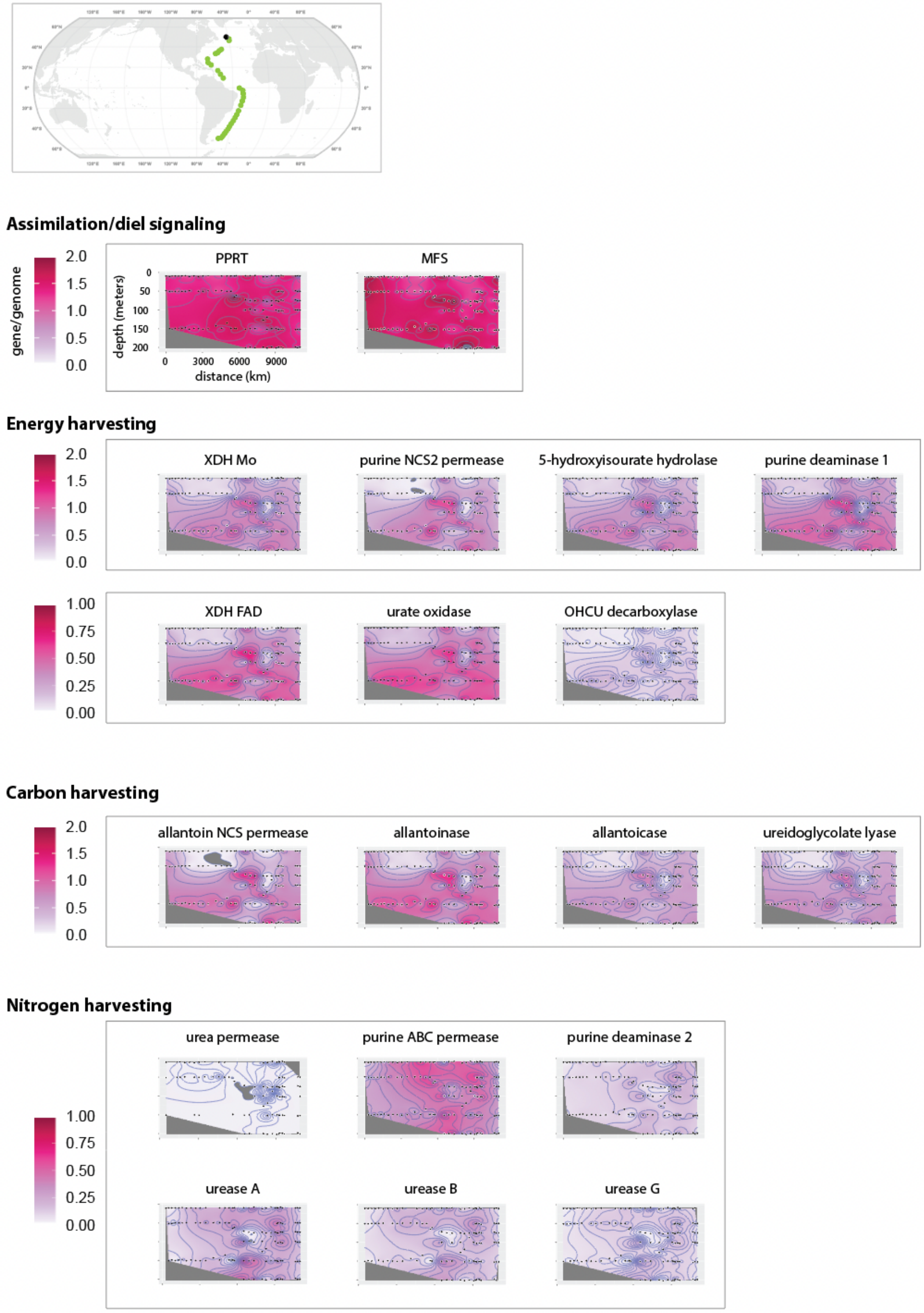

**Fig. S6. Depth based distribution of SAR11 purine usage genes along GA02 transect in Atlantic.** Black dot in top panel represents the first station of cruise transect.

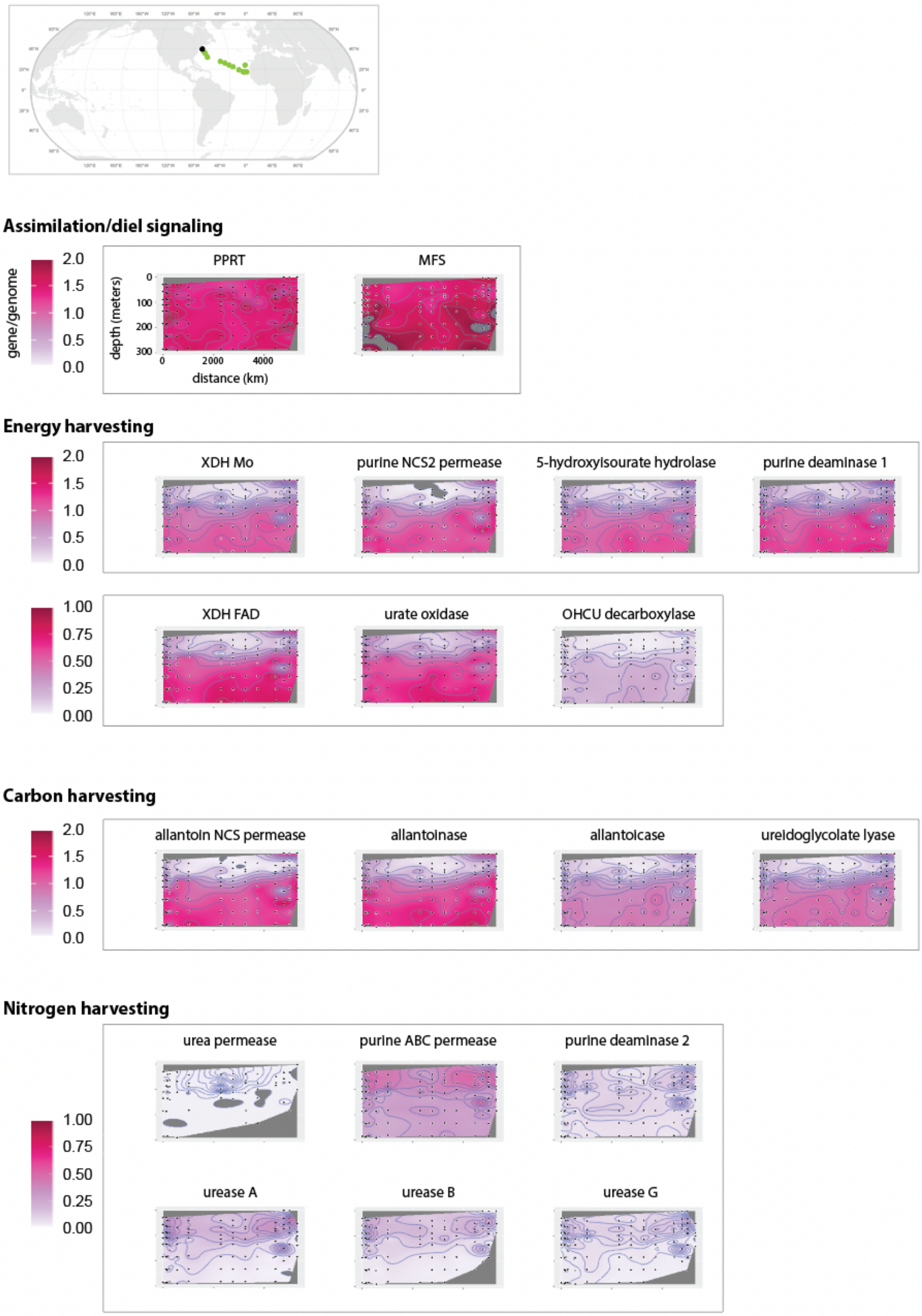

**Fig. S7. Depth based distribution of SAR11 purine usage genes along GA03 transect in North Atlantic.** Black dot in top panel represents the first station of cruise transect.

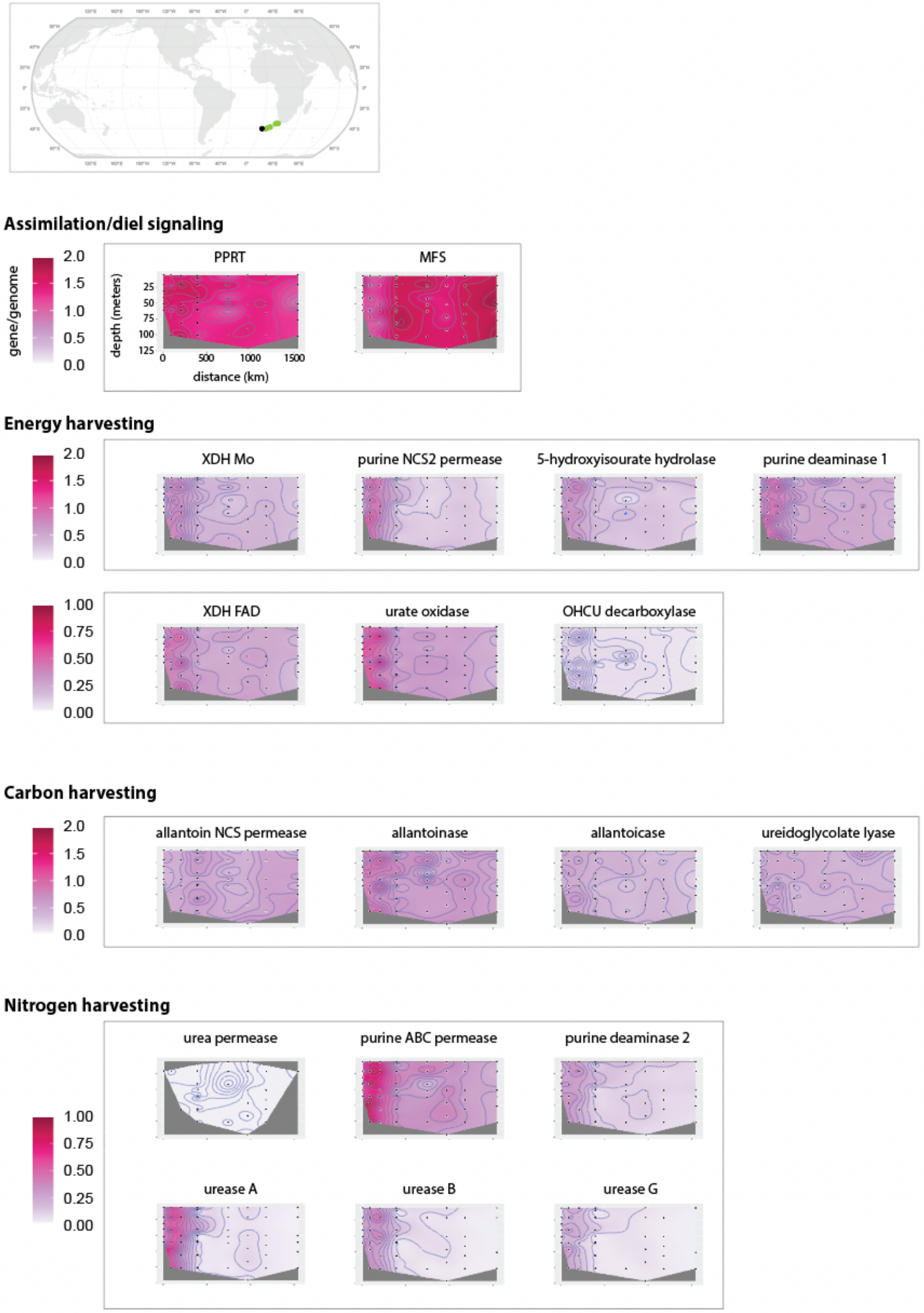

**Fig. S8. Depth based distribution of SAR11 purine usage genes for GA10 transect in South Atlantic.** Black dot in top panel represents the first station of cruise transect.

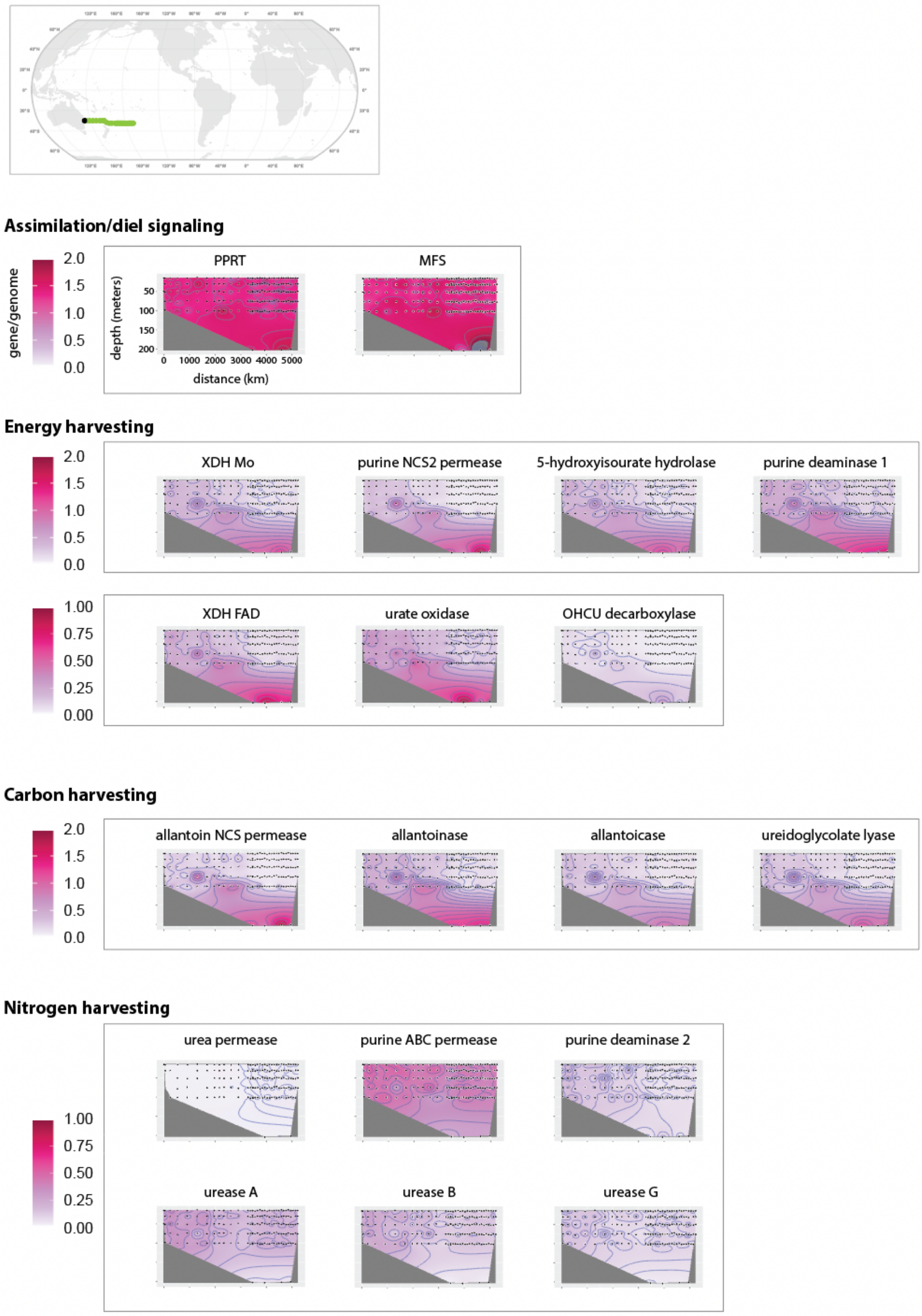

**Fig. S9. Depth based distribution of SAR11 purine usage genes for GP13 transect in South Pacific.** Black dot in top panel represents the first station of cruise transect.

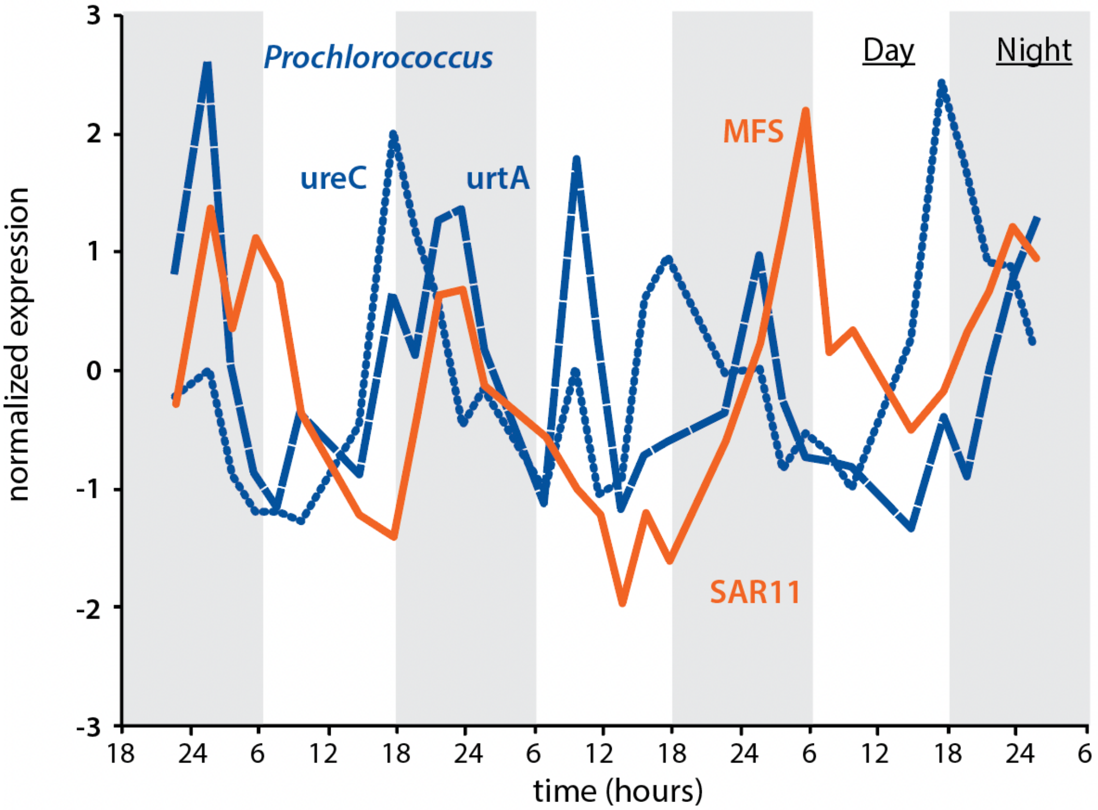

**Fig. S10 In situ expression of genes related to putative cross-feeding of urea between SAR11 and *Prochlorococcus*.** Normalized in situ expression of urease (ureC) and urea transport system (urtA) genes in *Prochlorococcus* (dark blue), and of SAR11 purine MFS permease (orange), in the North Pacific Ocean (data from Ref. 51).

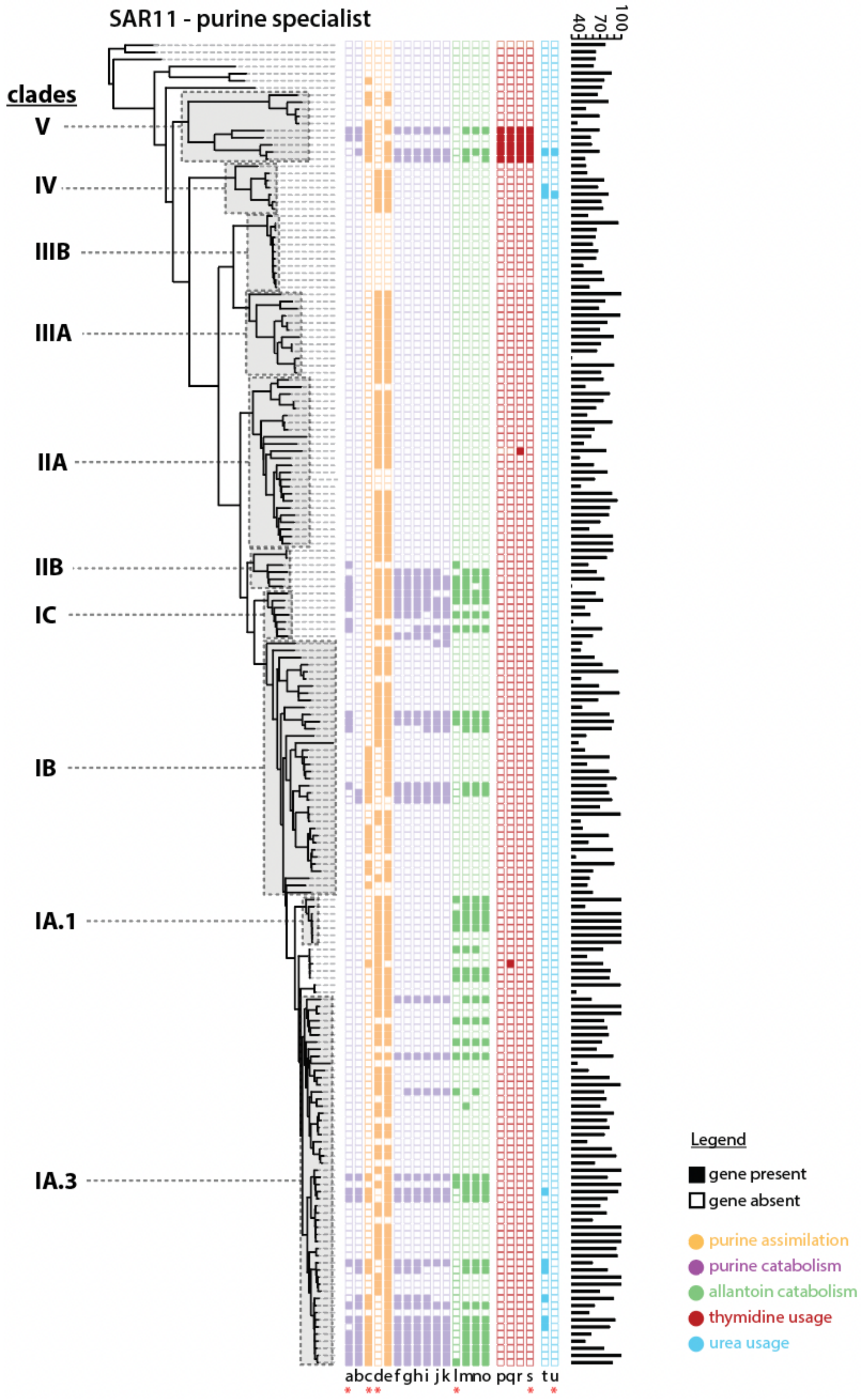

**Fig. S11. Diversity of purine usage strategies of SAR11 clades.** Rows in the phylometabolic diagrams represent the metabolic gene profiles within individual genomes in the tree. Transporter genes are indicated with pink asterisks. Colors represent different functions of pathways: purine assimilation genes are in yellow, purine catabolism genes in purple, allantoin catabolism genes in green, thymidine usage genes in dark red and urea assimilation genes in light blue. Genome completeness statistics are shown as histograms next to gene profiles. Genes: a = purine NCS2 permease, b = purine deaminase (tadA), c = purine ABC transporter permease, d = purine MFS permease, e = purine phosphoribosyltransferase (PPRT), f = urate oxidase (PuuD), g = purine deaminase (tadA), h = Xanthine dehydrogenase Mo-subunit (XDH_Mo) , i = xanthine dehydrogenase FAD-subunit (XDH_FAD) , j = 5-hydroxyisourate lyase (hiuH), k = 2-oxo-4-hydroxy-4-carboxy-5-ureidoimidazoline decarboxylase (OHCU decarboxylase, uraD), l = allantoin NCS1 permease, m = allantoicase (alc), n = ureidoglycolate lyase (ugl) , o = allantoinase (PuuE), p = uracil phosphoribosyltransferase, q = cytidine deaminase, r = thymidine kinase, s = pyrimidine ABC transporter, t = urease, u = urea ABC transporter

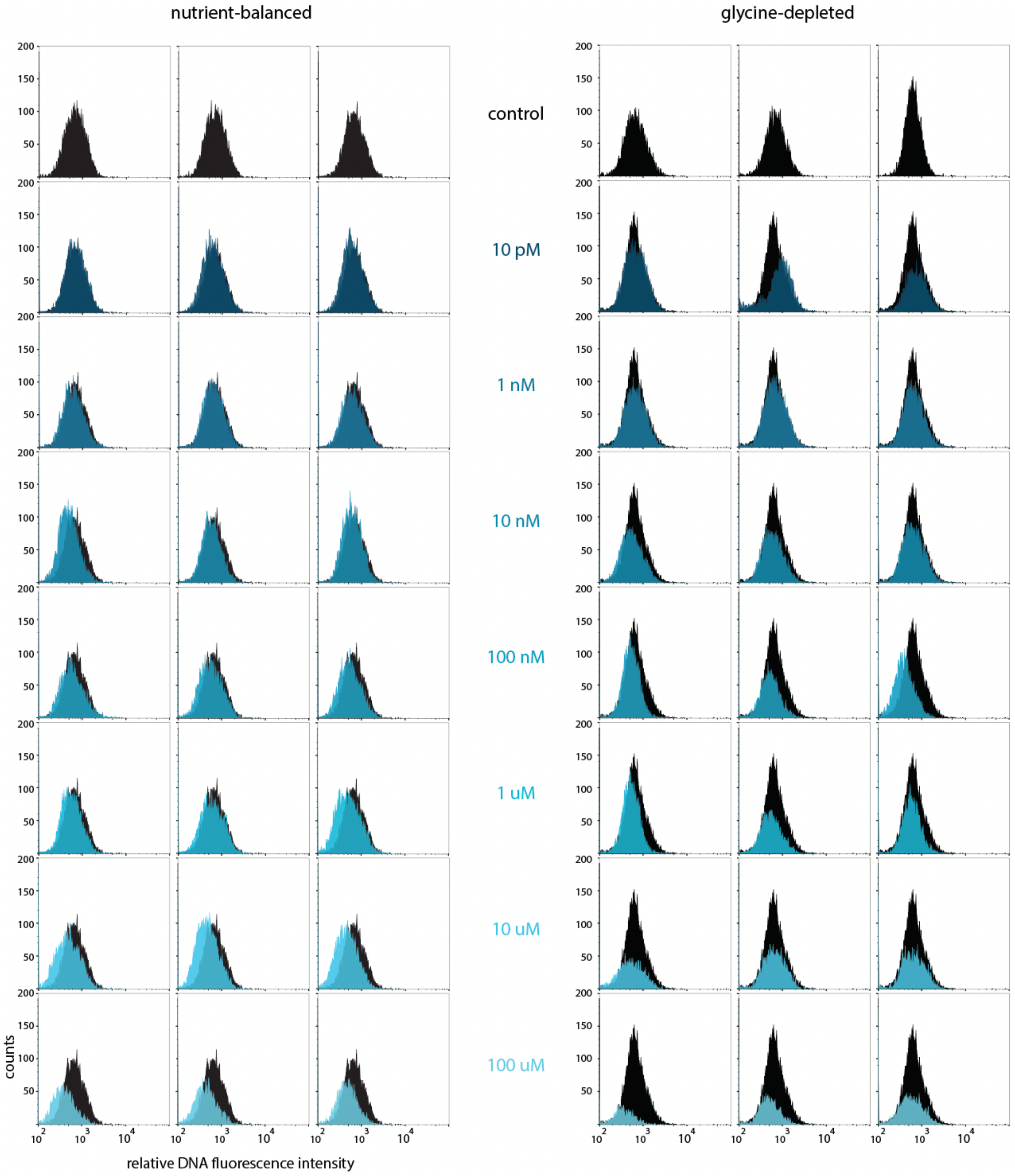

**Fig. S12. Distributions of relative DNA fluorescence during mid-exponential growth in SAR11 cultures amended with adenine.** Data shown are for day 7 of SAR11 cultures grown in both glycine-depleted and nutrient-balanced media that were amended with different concentrations of adenine (Fig. 7). Data from triplicate cultures treated with adenine at different concentrations is shown in different shades of blue as in Fig. 7 in the main text. Controls without adenine are shown as individual replicates in black at the top and were combined as superimposed histograms within the panels of each individual adenine-treated culture replicate to highlight the offset between cultures grown with and without adenine.

| Strain | **MIT9301** | | | | | | **MIT0801** | | **MIT9313** | | | |
|---|---|---|---|---|---|---|---|---|---|---|---|---|
| Genome size (MB) | 1.64 | | | | | | 1.93 | | 2.41 | | | |
| GC% | 31.3 | | | | | | 34.9 | | 50.7 | | | |
| Growth conditions* | *50, -P* | | *50, R* | | *10, R* | | *10, R* | | *10, R* | | *5, R* | |
| **A.** Extracellular levels (fg/cell) | | | | | | | | | | | | |
| | AVG | SD | AVG | SD | AVG | SD | AVG | SD | AVG | SD | AVG | SD |
| Thymidine | 1.9E-01 | 3.3E-02 | 1.5E-01 | 3.0E-02 | 1.4E-01 | 3.3E-02 | 1.6E-01 | 2.4E-02 | 6.2 E-02 | 2.3E-02 | 3.7E-02 | 5.4E-03 |
| Adenine** | 1.8E-03 | 5.2E-04 | (2.3E-04) | - | (2.3E-04) | - | ND | - | ND | - | ND | - |
| Guanine** | 1.5E-03 | 4.4E-04 | ND | - | ND | - | ND | - | ND | - | ND | - |
| Methylthioadenosine | 1.7E-03 | 3.6E-04 | ND | - | ND | - | ND | - | 3.8E-04 | 1.4E-04 | 2.3E-04 | 2.1E-04 |
| **B.** Extracellular levels normalized (molecules exuded / molecules incorporated) | | | | | | | | | | | | |
| | AVG | SD | AVG | SD | AVG | SD | AVG | SD | AVG | SD | AVG | SD |
| Thymidine | 4.2E-01 | 7.1E-02 | 3.3E-01 | 6.6E-02 | 3.1E-01 | 7.3E-02 | 3.2E-01 | 4.8E-02 | 1.3E-01 | 4.8E-02 | 7.7E-02 | 1.1E-02 |
| Adenine | 7.2E-03 | 2.1E-03 | (1.2E-03) | - | (9.1E-04) | - | ND | - | ND | - | ND | - |
| Guanine | 1.1E-02 | 3.4E-03 | ND | - | ND | - | ND | - | ND | - | ND | - |
| Methylthioadenosine | 3.0E-03 | 6.4E-04 | ND | - | ND | - | ND | - | 6.5E-04 | 2.4E-04 | 3.8E-04 | 3.6E-04 |
| **C.** Intracellular concentrations (μM) | | | | | | | | | | | | |
| | AVG | SD | AVG | SD | AVG | SD | AVG | SD | AVG | SD | AVG | SD |
| Thymidine | ND | - | ND | - | ND | - | ND | - | ND | - | ND | - |
| Adenine | 36.7 | 4.1 | ND | - | ND | - | ND | - | ND | - | ND | - |
| Guanine | ND | - | ND | - | ND | - | ND | - | ND | - | ND | - |
| Methylthioadenosine | 15.0 | 1.4 | 3.7 | 0.7 | ND | - | ND | - | 3.9 | 1.7 | ND | - |

**Table S1. Cumulative extracellular production levels and intracellular concentrations of thymidine, adenine, guanine and methylthioadenosine in three strains of *Prochlorococcus*.** ND indicates not detected. Values listed in (parentheses) were detected only in single replicates. *Growth conditions: numbers indicate light levels in units of mmol photons/$m^2$/s, *-P* indicates growth in semi-continuous phosphorus limitation, *R* indicates growth in nutrient replete batch cultures. **Uncorrected concentrations (Methods).

| | | $[NO_3^- + NO_2^-]$ | | $[PO_4^-]$ | | [Fe] | |
|---|---|---|---|---|---|---|---|
| | | $r_S$ | p | $r_S$ | p | $r_S$ | p |
| Nitrogen harvesting | Urea permease | -0.50 | 8.3E-28 | -0.37 | 3.6E-15 | -0.15 | 1.5E-03 |
| | Urease A | -0.50 | 2.2E-39 | -0.59 | 1.5E-58 | -0.21 | 2.7E-07 |
| | Urease B | -0.44 | 4.1E-28 | -0.53 | 1.9E-43 | -0.17 | 7.2E-05 |
| | Urease G | -0.43 | 4.3E-27 | -0.52 | 5.3E-42 | -0.21 | 4.4E-07 |
| | Purine ABC permease | -0.44 | 3.7E-31 | -0.46 | 3.5E-34 | -0.08 | 5.6E-02 |
| | Purine deaminase | -0.01 | 8.7E-01 | -0.11 | 4.5E-03 | -0.01 | 8.6E-01 |
| Assimilation/ Diel Signaling | Purine MFS permease | 0.20 | 4.8E-07 | 0.13 | 9.7E-04 | -0.08 | 5.9E-02 |
| | PPRT | 0.23 | 9.2E-09 | 0.12 | 3.9E-03 | 0.11 | 6.5E-03 |
| Energy harvesting | Purine NCS2 permease | 0.60 | 6.9E-62 | 0.45 | 1.6E-32 | 0.44 | 2.9E-31 |
| | 5-hydroxyisourate lyase | 0.61 | 5.6E-66 | 0.48 | 3.3E-37 | 0.50 | 7.4E-41 |
| | OHCU decarboxylase | 0.59 | 4.9E-60 | 0.48 | 2.6E-37 | 0.56 | 2.6E-51 |
| | Urate oxidase | 0.62 | 1.7E-68 | 0.49 | 4.6E-39 | 0.49 | 1.9E-38 |
| | Purine deaminase | 0.65 | 1.7E-74 | 0.50 | 6.7E-41 | 0.47 | 3.7E-35 |
| | XHD-Mo | 0.65 | 5.3E-76 | 0.51 | 4.3E-42 | 0.49 | 1.8E-38 |
| | XDH-FAD | 0.65 | 2.1E-76 | 0.53 | 3.0E-46 | 0.51 | 2.9E-42 |
| Carbon harvesting | Allantoinase | 0.71 | 2.5E-96 | 0.59 | 1.1E-58 | 0.42 | 8.8E-28 |
| | Allantoicase | 0.71 | 1.2E-95 | 0.61 | 3.0E-64 | 0.51 | 5.9E-43 |
| | Ureidoglycolate lyase | 0.72 | 2.8E-102 | 0.62 | 8.6E-67 | 0.49 | 9.5E-39 |
| | Allantoin NCS1 permease | 0.73 | 2.3E-106 | 0.60 | 2.5E-63 | 0.36 | 2.7E-20 |

**Table S2. Spearman rank correlation coefficients ($r_S$) between gene/genome frequencies of SAR11 purine usage genes in metagenomes and environmental nutrient concentrations.** Abbreviations: PPRT = purine phosphoribosyltransferase, OHCU decarboxylase = 2-oxo-4-hydroxy-4-carboxy-5-ureidoimidazoline decarboxylase, XDH = xanthine dehydrogenase.

| | | | Assimilation/ Diel Signaling | | | Energy harvesting* | | | | | | | Carbon harvesting | | | | Nitrogen harvesting | | | | |
|---|---|---|---|---|---|---|---|---|---|---|---|---|---|---|---|---|---|---|---|---|---|
| | # genomes | AVG completeness (%) | A | B | C | D | E | F | G | H | I | J | K | L | M | N | O | P | Q | R | S |
| *All* | 186 | 72 | 21 | 58 | 77 | 13 | 17 | 18 | 18 | 18 | 17 | 17 | 12 | 23 | 22 | 22 | 10 | 5 | 5 | 5 | 1 |
| **Clades** | | | | | | | | | | | | | | | | | | | | | |
| V | 10 | 61 | 80 | 0 | 70 | 20 | 30 | 30 | 30 | 30 | 30 | 30 | 0 | 30 | 20 | 30 | 30 | 10 | 10 | 10 | 10 |
| IV | 7 | 68 | 0 | 86 | 86 | 0 | 0 | 0 | 0 | 0 | 0 | 0 | 0 | 0 | 0 | 0 | 0 | 29 | 29 | 29 | 14 |
| IIIB | 11 | 65 | 0 | 0 | 0 | 0 | 0 | 0 | 0 | 0 | 0 | 0 | 0 | 0 | 0 | 0 | 0 | 0 | 0 | 0 | 0 |
| IIIA | 12 | 76 | 0 | 100 | 100 | 0 | 0 | 0 | 0 | 0 | 0 | 0 | 0 | 0 | 0 | 0 | 0 | 0 | 0 | 0 | 0 |
| IIA | 24 | 70 | 0 | 83 | 83 | 0 | 0 | 0 | 0 | 0 | 0 | 0 | 0 | 0 | 0 | 0 | 0 | 0 | 0 | 0 | 0 |
| IIB | 6 | 66 | 0 | 83 | 83 | 50 | 50 | 50 | 50 | 50 | 50 | 33 | 50 | 50 | 33 | 50 | 0 | 0 | 0 | 0 | 0 |
| IC | 7 | 58 | 0 | 86 | 86 | 71 | 71 | 71 | 86 | 71 | 57 | 71 | 57 | 57 | 57 | 57 | 0 | 0 | 0 | 0 | 0 |
| IB | 36 | 68 | 42 | 44 | 81 | 11 | 14 | 14 | 14 | 17 | 19 | 19 | 6 | 14 | 14 | 14 | 6 | 3 | 3 | 3 | 0 |
| IA.1 | 7 | 93 | 0 | 100 | 100 | 0 | 0 | 0 | 0 | 0 | 0 | 0 | 57 | 71 | 71 | 71 | 0 | 0 | 0 | 0 | 0 |
| IA.3 | 52 | 80 | 27 | 56 | 88 | 21 | 31 | 33 | 33 | 31 | 29 | 29 | 13 | 37 | 37 | 35 | 27 | 10 | 10 | 10 | 0 |

**Table S3. Frequency (%) of genes belonging to different purine usage functions in SAR11, broken down by clade.** Genes: A = Purine ABC transporter, B = Purine MFS transporter, C = purine phosphoribosyltransferase, D = Purine NCS2 permease, E = urate oxidase, F = purine deaminase, G = xanthine dehydrogenase Mo-subunit, H = xanthine dehydrogenase FAD-subunit, I = 5-hydroxyisourate lyase, J = OHCU decarboxylase, K = Allantoin NCS1 permease, L = Allantoicase, M = ureidoglycolate lyase, N = allantoinase, O = purine deaminase, P = urease A, Q = urease B, R = urease C, S = urea ABC transporter. Note: *Genes E-J are the core catabolic genes making up the energy harvesting portion of purine catabolism, with transporter gene D occasionally replaced by other transporters (Fig. 3).

| | MIT9313 | MIT0801 | MIT9301 | MED4 |
|---|---|---|---|---|
| Adenine phosphoribosyl transferase (apt) | PMT_0810 | EW15_1073 | P9301_14411 | PMM1122 |
| Methylthioadenosine phosphorylase (MTAP) | PMT_1900 | EW15_0393 | P9301_03251 | PMM0301 |
| 5'-nucleosidase (survival protein – SurE) | PMT_0366 | EW15_1747 | P9301_14561 | PMM1271 |
| FAMIN* | PMT_1235 | - | P9301_05541 | PMM0528 |
| Putative hypoxanthine-guanine phosphoribosyl transferase (hgpt) | - | EW15_0922 | P9301_10661 | PMM0985 |
| Uracil phosphoribosyl transferase (upp) | PMT_0558 | EW15_0864 | P9301_08361 | PMM0776 |
| Uridine kinase (udk) | - | EW15_1087 | P9301_10701 | PMM0989 |
| Cytosine deaminase (coda) | PMT_0327 | EW15_1782 | P9301_14901 | PMM1304 |
| Uracil phosphoribosyltransferase (pyrR) | PMT_1445 | - | P9301_16231 | PMM1433 |

**Table S4. KEGG gene IDs of possible genes involved in (deoxy)ribonucleotide recycling.** Entries marked with a (-) sign indicate no homolog of that gene was found in a given strain. Notes: *FAMIN is a highly multi-functional enzyme involved in many reactions within nucleotide metabolism (73)

| | dnaN | | dnaE | | dnaQ | | rpoC2 | | rpoC1 | | rpoB | |
|---|---|---|---|---|---|---|---|---|---|---|---|---|
| | $r_P$ | p | $r_P$ | p | $r_P$ | p | $r_P$ | p | $r_P$ | p | $r_P$ | p |
| apt | 0.062 | 0.767 | 0.025 | 0.904 | 0.063 | 0.764 | 0.488 | 0.013 | 0.518 | 0.008 | 0.647 | 4.78E-4 |
| MTAP | 0.531 | 0.006 | 0.367 | 0.071 | 0.329 | 0.108 | 0.841 | 1.39E-7 | 0.822 | 4.79E-7 | 0.745 | 1.93E-5 |
| SurE | -0.045 | 0.831 | -0.232 | 0.264 | -0.262 | 0.021 | 0.585 | 0.002 | 0.725 | 4.09E-5 | 0.601 | 0.001 |
| FAMIN | -0.439 | 0.028 | -0.564 | 0.003 | -0.568 | 0.003 | 0.097 | 0.643 | 0.338 | 0.098 | 0.199 | 0.339 |
| hgpt | 0.697 | 1.08E-4 | 0.653 | 3.99E-4 | 0.628 | 7.8E-4 | 0.464 | 0.019 | 0.379 | 0.062 | 0.302 | 0.142 |
| upp | -0.124 | 0.554 | -0.328 | 0.109 | -0.381 | 0.060 | 0.440 | 0.028 | 0.634 | 6.63E-4 | 0.370 | 0.069 |
| udk | 0.350 | 0.086 | 0.317 | 0.123 | 0.303 | 0.141 | 0.422 | 0.036 | 0.380 | 0.061 | 0.384 | 0.058 |
| CodA | -0.422 | 0.035 | -0.590 | 0.002 | -0.612 | 0.001 | 0.154 | 0.462 | 0.400 | 0.048 | 0.184 | 0.380 |
| pyrR | 0.259 | 0.212 | 0.088 | 0.674 | 0.053 | 0.801 | 0.733 | 3.07E-5 | 0.796 | 1.95E-6 | 0.711 | 6.71E-5 |

**Table S5. Pearson correlation coefficients ($r_P$) and significance (p) of temporal expression profiles of possible genes involved in (deoxy)ribonucleotide recycling and DNA/RNA polymerase genes in *Prochlorococcus*.** Data is from *Prochlorococcus* MED4 and was obtained from a study in which growth was synchronized to diurnal L:D conditions (29).

| rP | Nitrogen harvesting | | | | | Assimilation | | Energy harvesting | | | | | | | Carbon harvesting | | | |
|---|---|---|---|---|---|---|---|---|---|---|---|---|---|---|---|---|---|---|
| | A | B | C | D | E | F | G | H | I | J | K | L | M | N | O | P | Q | R |
| A: Purine deaminase 2 | 1 | 0.67 | 0.54 | 0.58 | 0.55 | -0.48 | 0.17 | 0.07 | 0.13 | 0.14 | 0.14 | 0.23 | 0.23 | 0.14 | -0.11 | 0.21 | 0.11 | 0.15 |
| B: ABC permease | | 1 | 0.78 | 0.76 | 0.76 | -0.40 | 0.07 | -0.26 | -0.27 | -0.32 | -0.28 | -0.23 | -0.29 | -0.28 | -0.48 | -0.33 | -0.33 | -0.38 |
| C: Urease A | | | 1 | 0.93 | 0.94 | -0.21 | 0.15 | -0.25 | -0.25 | -0.34 | -0.28 | -0.23 | -0.37 | -0.27 | -0.47 | -0.39 | -0.32 | -0.42 |
| D: Urease B | | | | 1 | 0.90 | -0.30 | 0.11 | -0.28 | -0.26 | -0.32 | -0.28 | -0.21 | -0.31 | -0.26 | -0.49 | -0.37 | -0.32 | -0.40 |
| E: Urease C | | | | | 1 | -0.27 | 0.10 | -0.26 | -0.26 | -0.34 | -0.28 | -0.24 | -0.34 | -0.28 | -0.48 | -0.37 | -0.33 | -0.40 |
| F: MFS permease | | | | | | 1 | 0.50 | 0.48 | 0.37 | 0.22 | 0.33 | 0.17 | -0.11 | 0.28 | 0.56 | 0.11 | 0.41 | 0.20 |
| G: PPRT | | | | | | | 1 | 0.52 | 0.49 | 0.39 | 0.47 | 0.43 | 0.15 | 0.43 | 0.45 | 0.29 | 0.49 | 0.34 |
| H: NCS2 permease | | | | | | | | 1 | 0.95 | 0.88 | 0.96 | 0.85 | 0.65 | 0.92 | 0.94 | 0.83 | 0.95 | 0.87 |
| I: Purine deaminase 1 | | | | | | | | | 1 | 0.96 | 0.99 | 0.95 | 0.76 | 0.97 | 0.92 | 0.89 | 0.97 | 0.92 |
| J: XDH FAD | | | | | | | | | | 1 | 0.97 | 0.97 | 0.89 | 0.97 | 0.86 | 0.94 | 0.94 | 0.95 |
| K: XDH Mo | | | | | | | | | | | 1 | 0.95 | 0.81 | 0.98 | 0.92 | 0.93 | 0.97 | 0.95 |
| L: Urate oxidase | | | | | | | | | | | | 1 | 0.86 | 0.96 | 0.82 | 0.91 | 0.92 | 0.92 |
| M: OHCU decarboxylase | | | | | | | | | | | | | 1 | 0.82 | 0.64 | 0.93 | 0.73 | 0.89 |
| N: 5-hydroxyisurate lyase | | | | | | | | | | | | | | 1 | 0.88 | 0.91 | 0.95 | 0.93 |
| O: Allantoin permease | | | | | | | | | | | | | | | 1 | 0.84 | 0.95 | 0.89 |
| P: Allantoicase | | | | | | | | | | | | | | | | 1 | 0.89 | 0.98 |
| Q: Allantoinase | | | | | | | | | | | | | | | | | 1 | 0.93 |
| R: Ureidoglycolate lyase | | | | | | | | | | | | | | | | | | 1 |
| p | | | | | | | | | | | | | | | | | | |
| A: Purine deaminase 2 | 1 | 5.8E-90 | 1.1E-52 | 4.9E-64 | 7.6E-55 | 6.6E-40 | 6.3E-06 | 5.7E-02 | 9.7E-04 | 2.9E-04 | 1.4E-04 | 5.3E-10 | 9.2E-10 | 1.7E-04 | 4.8E-03 | 2.9E-08 | 4.1E-03 | 1.1E-04 |
| B: ABC permease | | 1 | 3.6E-144 | 8.0E-132 | 3.1E-129 | 1.0E-27 | 6.5E-02 | 3.5E-12 | 2.4E-13 | 4.1E-18 | 3.9E-14 | 1.1E-09 | 8.3E-15 | 3.4E-14 | 3.2E-41 | 6.4E-19 | 3.7E-19 | 3.3E-25 |
| C: Urease A | | | 1 | 0 | 0 | 4.1E-08 | 5.3E-05 | 5.1E-11 | 1.8E-11 | 1.5E-19 | 1.1E-13 | 1.8E-09 | 6.8E-24 | 2.6E-13 | 4.7E-39 | 5.0E-27 | 1.5E-17 | 5.4E-31 |
| D: Urease B | | | | 1 | 5.1E-253 | 6.0E-16 | 3.3E-03 | 7.6E-14 | 5.5E-12 | 2.7E-17 | 5.5E-14 | 2.6E-08 | 6.1E-17 | 2.4E-12 | 3.1E-43 | 7.1E-24 | 3.1E-18 | 4.3E-28 |
| E: Urease C | | | | | 1 | 3.4E-13 | 9.6E-03 | 2.6E-12 | 1.8E-12 | 1.4E-19 | 4.0E-14 | 3.1E-10 | 1.2E-19 | 5.8E-14 | 5.2E-40 | 5.1E-24 | 8.6E-19 | 4.2E-28 |
| F: MFS permease | | | | | | 1 | 2.4E-44 | 2.0E-40 | 2.8E-23 | 6.5E-09 | 2.4E-19 | 7.6E-06 | 3.3E-03 | 7.1E-14 | 7.4E-57 | 3.3E-03 | 1.6E-29 | 9.5E-08 |
| G: PPRT | | | | | | | 1 | 4.7E-48 | 1.8E-43 | 9.3E-27 | 3.7E-38 | 6.9E-32 | 5.6E-05 | 1.6E-32 | 3.9E-35 | 4.3E-15 | 5.6E-43 | 6.8E-20 |
| H: NCS2 permease | | | | | | | | 1 | 0 | 9.8E-226 | 0 | 2.7E-194 | 4.3E-84 | 5.0E-276 | 0 | 9.1E-175 | 0 | 1.7E-215 |
| I: Purine deaminase 1 | | | | | | | | | 1 | 0 | 0 | 0 | 1.9E-132 | 0 | 2.9E-274 | 3.1E-232 | 0 | 1.4E-282 |
| J: XDH FAD | | | | | | | | | | 1 | 0 | 0 | 2.1E-235 | 0 | 7.6E-207 | 0 | 0 | 0 |
| K: XDH Mo | | | | | | | | | | | 1 | 0 | 1.3E-161 | 0 | 9.6E-283 | 2.4E-295 | 0 | 0 |
| L: Urate oxidase | | | | | | | | | | | | 1 | 2.2E-201 | 0 | 3.1E-167 | 3.0E-266 | 3.2E-289 | 6.2E-285 |
| M: OHCU decarboxylase | | | | | | | | | | | | | 1 | 2.0E-171 | 2.6E-81 | 1.1E-296 | 8.7E-114 | 1.3E-230 |
| N: 5-hydroxyisurate lyase | | | | | | | | | | | | | | 1 | 5.1E-226 | 1.0E-259 | 0 | 1.9E-303 |
| O: Allantoin permease | | | | | | | | | | | | | | | 1 | 5.4E-181 | 0 | 1.1E-234 |
| P: Allantoicase | | | | | | | | | | | | | | | | 1 | 1.3E-231 | 0 |
| Q: Allantoinase | | | | | | | | | | | | | | | | | 1 | 1.4E-297 |
| R: Ureidoglycolate lyase | | | | | | | | | | | | | | | | | | 1 |

**Table S6. Pearson correlation coefficients ($r_P$) with significance values (p) among gene/genome frequencies of SAR11 purine usage genes belonging to different pathway functions.** Genes (labeled A-R) in different functional categories listed at top are defined on the left side of the table.
